# Logic-Gated Fluorogenic RNA Reporters for Multiplexed Live-Cell Imaging

**DOI:** 10.64898/2026.08.16.745110

**Authors:** Sima Khajouei, Ali Ebrahimi Darsinouei, Ru Zheng, Jeffrey Chen, Qinge Liu, Zhaolin Xue, Mingxu You

**Author notes:** These authors contribute equally (Sima Khajouei, Ali Ebrahimi Darsinouei). Corresponding author E-mail address (M. You).

## Abstract

Multiplexed imaging of biomolecular networks in living cells is limited by the small number of spectral separable fluorophores and the need to monitor dynamic processes in real time. Here, we present logicFRIES, a fluorogenic RNA (FR)-based platform that enables eight-plex live-cell imaging through programmable, logic-gated activation coupled with sequential fluorescence imaging. By integrating small-molecule-binding RNA aptamers into dye-activating fluorogenic RNAs, we engineered trigger-responsive FR reporters. This design implements an AND-gated mechanism in which fluorescence activation requires both a cognate trigger molecule and its corresponding fluorogenic dye, thereby expanding multiplexing capacity without adding new fluorophores. Using three membrane-permeable triggers, tetracycline, ASP2905, and guanine, we generated three distinct trigger-defined activation states for each engineered Broccoli and Pepper FR. Combined with orthogonal Corn/DFHO and DNB/TMR-DN reporter pairs, logicFRIES supports eight-plex imaging through sequential trigger/dye addition, imaging, and wash-based stripping cycles. We demonstrate robust, specific, and reversible fluorescence switching of these multiplexed FR reporters in living HEK293T and SKBR3 cells. Overall, logicFRIES extends live-cell imaging beyond conventional spectral limitations and provides a modular foundation for potentially developing multiplexed sensors targeting endogenous RNAs, proteins, and small molecules in complex cellular systems.

## Introduction

Cellular functions arise from intricate networks of interacting biomolecules, whose spatial organization and dynamic coordination regulate cell architecture, signaling fidelity, and overall phenotype [1–3]. Elucidating how these components act together in living cells is critical for understanding normal physiology and disease. Achieving this goal requires imaging strategies that can detect multiple molecular targets *in situ*, in real time, and within their native cellular environment. As a widely adopted platform for live-cell imaging, fluorescence microscopy combines sensitive signal readout with high temporal resolution and minimal disruption to biological processes. However, its multiplexing capacity is fundamentally constrained by the broad emission spectra of fluorophores and their significant spectral overlap, which typically limits simultaneous detection to only a few targets [4–6] Consequently, capturing the coordinated behavior of multiple biomolecules, especially RNAs and proteins, within living cells remains a major technical challenge.

In fixed and permeabilized specimens, these constraints have been mitigated through multiplexed spatial profiling techniques. The advent of transcriptome-scale RNA imaging methods such as MERFISH, seqFISH, STARmap, and FISSEQ have enabled large-scale RNA mapping *in situ* with high spatial resolution [7–10]. While these methods have transformed spatial molecular profiling, their reliance on fixation and permeabilization precludes direct observation of dynamic processes in living cells. To overcome this limitation, recent work has advanced live-cell imaging modalities that enable tracking of biomolecular dynamics under native conditions, including especially genetically encoded fluorescent protein tags, CRISPR-derived imaging systems, and fluorogenic RNA aptamers [11–16]. Despite these progresses, most current live-cell imaging approaches remain limited to a simultaneous detection of only a small number of targets, often ≤4 for RNAs [4]. This restricted multiplexing capacity is insufficient to capture the coordinated activity of complex molecular networks, underscoring the need for new approaches that expand live-cell multiplexing while maintaining high spatial and temporal performance.

Herein, we developed a multiplex imaging platform, **logicFRIES**, based on fluorogenic RNA aptamers (**FR**s) that enables visualization of at least eight targets in living cells. FRs are single-stranded RNA aptamers identified via systematic evolution of ligands by exponential enrichment (SELEX). These genetically encodable structured RNAs selectively bind otherwise nonfluorescent small-molecule dyes and activate fluorescence only upon complex formation (Scheme 1A) [17,18]. LogicFRIES is built on our previously introduced sequential Fluogenic RNA Imaging-Enabled Sensor (seqFRIES) platform [16], in which FRs are engineered to report on endogenous targets and generate fluorescence sequentially upon the addition of their cognate membrane-permeable dyes. In this framework, four orthogonal FRs, i.e., Broccoli, Pepper, DNB, and Corn, are genetically encoded in cells, whereas the corresponding dyes (DFHBI-1T, HBC620, TMR-DN, and DFHO) are supplied exogenously, imaged, and then removed to extinguish fluorescence. Iterative cycles of dye addition, imaging, and washout enable sequential multiplexed detection in living cells using these noncovalent FR/dye complexes.

In logicFRIES, live-cell multiplexing capacity is further increased through conditional, logic-gated activation. Specifically, we embed a trigger-binding RNA aptamer within the stem region of an FR such that the dye-binding pocket assembles only when both the cognate small-molecule trigger and the corresponding fluorogenic dye are present, thereby implementing a logical AND gate (Scheme 1B–D). This AND-gated design couples fluorogenic activation to two independent inputs (trigger binding and dye binding), substantially increasing the number of distinguishable targets without introducing additional fluorophores. Using Broccoli and Pepper as examples, we demonstrate logicFRIES by integrating, for each FR, three distinct trigger-binding RNA aptamers that respond to membrane-permeable, nontoxic triggers (i.e., tetracycline, ASP2905, and guanine). This configuration yields three trigger-activated states per FR, producing six distinguishable signals from Broccoli- and Pepper-based reporters alone (Scheme 1B,C). When these six logic-gated reporters are combined with the two fully orthogonal FR/dye pairs established in seqFRIES (DNB/TMR-DN and Corn/DFHO), the platform supports eight-plex imaging via sequential addition of the appropriate triggers and dyes, followed by imaging and washing steps to reversibly extinguish fluorescence (Scheme 1E). By expanding the toolset of orthogonal fluorogenic RNAs and corresponding trigger molecules, this platform can be readily extended and programmed to support imaging of progressively larger numbers of targets in living cells.

## Results and Discussion

### Design and optimization of trigger-activated fluorogenic RNA reporters

To establish a generalizable strategy for AND-gated logic reporter construction and target imaging, we first focused on fusing a trigger-binding aptamer to an FR, similar to prior designs of allosteric RNA sensors [19–23]. We selected three previously reported cell-permeable triggers, guanine, ASP2905, and tetracycline, for which cognate RNA aptamers are available and that have been reported to be compatible with mammalian cell systems [24–27]. We prioritized the Broccoli/DFHBI-1T and Pepper/HBC620 reporter pairs because (1) allosteric sensors based on these FRs have been previously reported, (2) their architectures are well characterized and relatively straightforward to engineer, and (3) they provide bright fluorescence signals in living cells.

For Pepper-based logic reporters, we engineered a new Pepper-ASP2905 construct and adopted previously reported Pepper-guanine and Pepper-tetracycline sensors [28]. These trigger-activated Pepper logic reporters were designed as modular RNA devices comprising three functional elements: the Pepper fluorogenic RNA, a ligand-specific RNA aptamer that recognizes the trigger, and a transducer domain that couples trigger binding to the folding of Pepper. To maintain Pepper functionality, the trigger aptamer and transducer were inserted into a structurally essential region of the RNA. Prior mutational analyses identified the P1 stem as critical for folding and fluorescence of circularly permuted Pepper (Figure S1A) [28]. Accordingly, we selected this P1 stem as the integration site and engineered a panel of Pepper-ASP2905 constructs (Figure S1B).

In these designs, the transducer domain remains largely unpaired in the absence of the trigger, thereby disfavoring formation of the folded Pepper. Trigger binding stabilizes the aptamer and promotes hybridization of the transducer strands, restoring the Pepper structure and activating fluorescence. Following *in silico* design, we constructed and evaluated four transducer variants (PA1–PA4) (Figure S1C). Among them, PA2 and PA4 produced the greatest fluorescence enhancements (∼2.0-fold). After further optimization of RNA, dye, and Mg^2+^ concentrations, the construct incorporating transducer 2 (named as **PA**) achieved ∼7.4-fold activation (Figures 1A and S1D) and was selected for subsequent studies. Consistent with prior reports, the adopted Pepper-tetracycline (**PT**) and Pepper-guanine (**PG**) reporters exhibited ∼3.9-fold and ∼2.0-fold fluorescence activation *in vitro*, respectively (Figure 1B,C).

**Figure 1.**
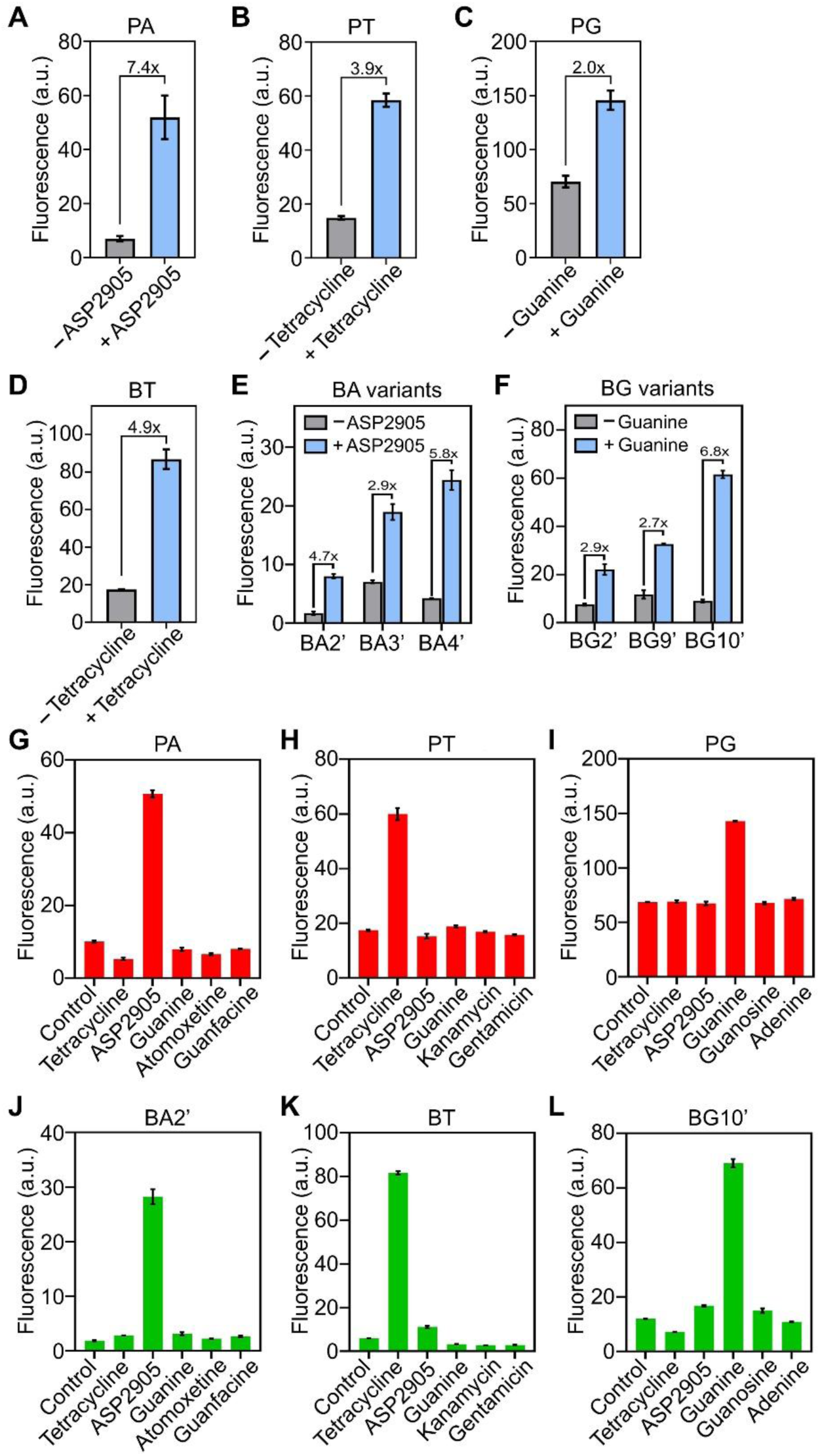
*In vitro* characterization of trigger-activated logicFRIES reporters. (**A–C**) Trigger-dependent fluorescence activation of the optimized ASP2905-, tetracycline-, and guanine-responsive Pepper reporters (PA, PT, and PG) following incubation with 5 µM ASP2905, 100 µM tetracycline, or 100 µM guanine, respectively, or in the absence of trigger. Reactions contained 1 µM RNA and 1 µM HBC620 dye. (**D–F**) Trigger-dependent fluorescence activation of the optimized ASP2905-, tetracycline-, and guanine-responsive Broccoli reporters (BA variants, BT, and BG variants) following incubation with 5 µM ASP2905, 100 µM tetracycline, or 100 µM guanine, respectively, or in the absence of trigger. Reactions contained 1 µM RNA and 20 µM DFHBI-1T dye. (**G–I**) Selectivity and trigger orthogonality of Pepper reporters assessed against noncognate triggers and structurally related control molecules (ASP2905, atomoxetine, guanfacine; tetracycline, kanamycin, gentamicin; guanine, guanosine, adenine). Trigger concentrations were 1 µM ASP2905/controls, 100 µM tetracycline/controls, and 10 µM guanine/controls; no-trigger samples served as the negative “control”. Reactions contained 1 µM RNA and 1 µM HBC620 dye. (**J–L**) Selectivity and trigger orthogonality of Broccoli reporters evaluated under the same conditions as in (G–I), using 1 µM RNA and 20 µM DFHBI-1T dye. All data are reported as mean ± standard deviation (SD) from at least three independent experiments and are normalized to the fluorescence of the corresponding positive controls (Pepper/HBC620 or Broccoli/DFHBI-1T) measured under matched conditions.

For Broccoli, we started from a reported Broccoli-tetracycline allosteric architecture and introduced minor structural modifications by extending the 5′ and 3′ termini to improve folding and fluorescence enhancement. In the presence of both tetracycline and DFHBI-1T, ∼4.9-fold activation was observed (**BT**, Figure 1D). In contrast, our initial allosteric designs for Broccoli-ASP2905 (BA1–BA6) and Broccoli-guanine (BG1–BG6) exhibited either elevated background fluorescence or insufficient trigger-dependent signal activation (Figure S2A,B). We therefore implemented an alternative RNA nanodevice framework developed by the Jaffrey group [29]. In this approach, an F30 RNA scaffold replaces the conventional duplex transducer and provides a three-way junction architecture that supports ligand-dependent control of Broccoli folding (Figure S2C). Guided by thermodynamic modeling, we fixed the Broccoli output stem while systematically varied the length and sequence of the input stem appended to the trigger-binding aptamer, enabling rational tuning of trigger coupling. Using this scaffold, multiple Broccoli-ASP2905 variants showed robust activation. Transducers 2’, 3’, and 4’ (BA2’–BA4’) exhibited 2.9–5.8-fold fluorescence enhancements (Figures 1E and S2D) and were advanced for cellular evaluation. For Broccoli-guanine reporters, transducers 2’, 9’, and 10’ (BG2’, BG9’, BG10’) yielded the strongest guanine-dependent responses (2.7–6.8-fold) and were similarly prioritized for further cellular studies (Figures 1F and S2E).

We next assessed the selectivity and trigger orthogonality of each logic reporter. Orthogonality is essential for logicFRIES, as each engineered logic FR must respond exclusively to its cognate trigger. As demonstrated in the subsequent cellular assessment, Broccoli-ASP2905 BA2’ and Broccoli-guanine BG10’ were selected for evaluation. For both Broccoli- and Pepper-based reporters, we observed negligible cross-reactivity among tetracycline, ASP2905, and guanine, confirming robust trigger orthogonality (Figure 1G–L). To evaluate specificity, we tested structurally related, noncognate small molecules as controls. For tetracycline-responsive reporters, we examined the aminoglycosides kanamycin and gentamicin. For ASP2905-responsive reporters, we used atomoxetine and guanfacine. For guanine-responsive reporters, we evaluated the purine analogs adenine and guanosine. In each case, the reporters responded selectively to its cognate trigger, while exhibiting minimal fluorescence activation in the presence of the noncognate controls (Figure 1G–L). Collectively, these six trigger-responsive FR constructs showed robust *in vitro* performance, providing a solid foundation for subsequent evaluation in living cells.

### Cellular performance and orthogonality of trigger-gated fluorogenic RNA reporters

To determine whether the engineered reporters function in living human cells, we evaluated these trigger-activated FRs in HEK293T cells. To promote robust intracellular RNA accumulation, each construct was expressed as a circular RNA using the Tornado system in the pAV-U6+27 vector [30]. For Pepper-based trigger-activated reporters, cells were imaged in the presence of 1 µM HBC620 together with the cognate triggers (30 µM tetracycline, 5 µM ASP2905, or 250 µM guanine). Co-incubation with HBC620 and the corresponding trigger resulted in strong fluorescence signals (Figure 2A), with an ∼8.8-, 5.6-, and 9.6-fold enhancement for PT, PA, and PG, respectively, indicating efficient activation in living cells.

**Figure 2.**
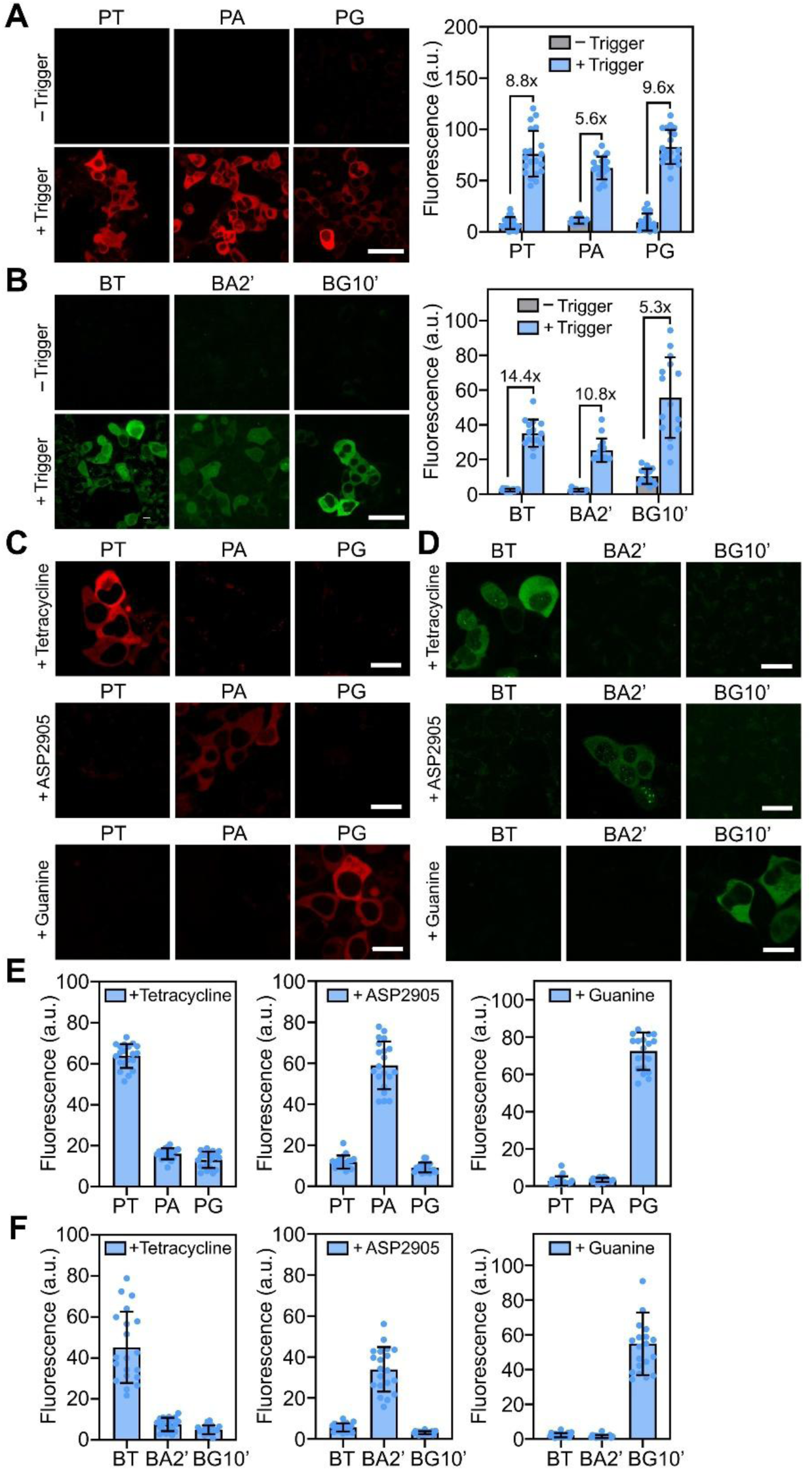
Cellular characterization and orthogonality of trigger-activated FR reporters. (**A**) Fluorescence imaging of HEK293T cells expressing trigger-activated Pepper constructs from the pAV-U6+27 vector. Cells were incubated with 1 μM HBC620 dye and the corresponding cognate triggers (30 μM tetracycline for PT, 5 μM ASP2905 for PA, and 250 μM guanine for PG) at 37°C for ∼30 min. Scale bar, 50 μm. Cellular fluorescence intensities were quantified from ∼20 individual cells per condition. Data are presented as mean ± SD from images acquired in at least three independent experiments and normalized to Pepper positive controls after incubation with 1 μM HBC620. (**B**) Fluorescence imaging of HEK293T cells expressing circular trigger-activated Broccoli constructs from the pAV-U6+27 vector after incubation with 40 μM DFHBI-1T dye and the corresponding triggers (30 μM tetracycline for BT, 5 μM ASP2905 for BA2’, and 250 μM guanine for BG10’) at 37°C for ∼30 min. Scale bar, 50 μm. Intracellular fluorescence intensities were quantified from ∼15 individual cells per condition. Data are presented as mean ± SD from images acquired in at least three independent experiments and normalized to Broccoli positive controls after incubation with 40 μM DFHBI-1T. (**C**) Orthogonality of trigger-responsive Pepper reporters as tested in HEK293T cells. PT, PA, and PG reporters were incubated with 1 μM HBC620 and each trigger (30 μM tetracycline, 5 μM ASP2905, or 250 μM guanine) in separate wells. Scale bar, 20 μm. (**D**) Orthogonality of Broccoli-based BT, BA2’, and BG10’ reporters after incubating with 40 μM DFHBI-1T and individual trigger molecules (30 μM tetracycline, 5 μM ASP2905, or 250 μM guanine). Scale bar, 20 μm. (**E**, **F**) Quantification of trigger-activated FR reporters in response to cognate and non-cognate triggers, measured from ∼20 individual cells per condition. Data are presented as mean ± SD from images acquired in at least three independent experiments and normalized to the corresponding Pepper/HBC620 or Broccoli/DFHBI-1T positive controls.

For Broccoli-based reporters, the same trigger concentrations were applied. The Broccoli-tetracycline construct exhibited a robust fluorescence response, yielding a ∼14-fold enhancement by confocal microscopy (Figure 2B). For Broccoli-ASP2905, guided by *in vitro* screening, BA2’, BA3’, and BA4’ constructs were evaluated (Figure S3). The **BA2’** construct produced the strongest response, with a ∼10-fold increase in fluorescence (Figure 2B). For Broccoli-guanine, BG2’, BG9’, and BG10’ constructs were tested. Among these, **BG10’** demonstrated the highest activation, with a ∼5-fold fluorescence enhancement (Figures 2B and S3). Collectively, these results confirm that the designed reporters can be effectively activated by their respective triggers and are capable of generating detectable fluorescence in living cells, highlighting their potential for multiplexed RNA imaging applications.

We next tested orthogonality of the six trigger-responsive reporters in HEK293T cells. Building on our prior demonstration that Broccoli/DFHBI-1T and Pepper/HBC620 function as an orthogonal reporter pair, we evaluated cross-reactivity among the trigger molecules. Pepper-based reporters were incubated with HBC620 and each trigger (ASP2905, guanine, or tetracycline) in separate wells (Figure 2C), while Broccoli-based reporters were incubated with 40 μM DFHBI-1T and the same set of triggers (Figure 2D). Quantitative analysis showed that each cognate RNA-dye-trigger combination produced fluorescence signals exceeding those of noncognate combinations by ∼4–25-fold for Pepper reporters and ∼6–40-fold for Broccoli reporters (Figure 2E,F), confirming strong orthogonality in living cells. Together, these data indicate that Broccoli- and Pepper-based logic reporters retain robust orthogonality in the presence of the corresponding triggers, supporting their use in multiplexed live-cell imaging.

### Live-cell kinetics of trigger-activated logicFRIES reporters

To meet the requirements of rapid sequential imaging, we next quantified the fluorescence activation and deactivation kinetics of each logicFRIES reporter in living cells. For activation measurements, cells were co-incubated with the appropriate dye and cognate trigger, and fluorescence intensity was monitored over 30 min. As shown in Figure 3A, trigger-activated Broccoli reporters exhibited rapid fluorescence activation in mammalian cells: ∼50% of maximal cellular fluorescence was reached within ∼11 min (BT), ∼8 min (BA2’), and ∼25 min (BG10’) following treatment with DFHBI-1T and the corresponding trigger (Figure 3C–E), indicating a fast and efficient trigger-dependent activation. Notably, although BG10′ displays slightly slower kinetics, robust cellular fluorescence signals are still readily detectable within the first 5 min of incubation.

**Figure 3.**
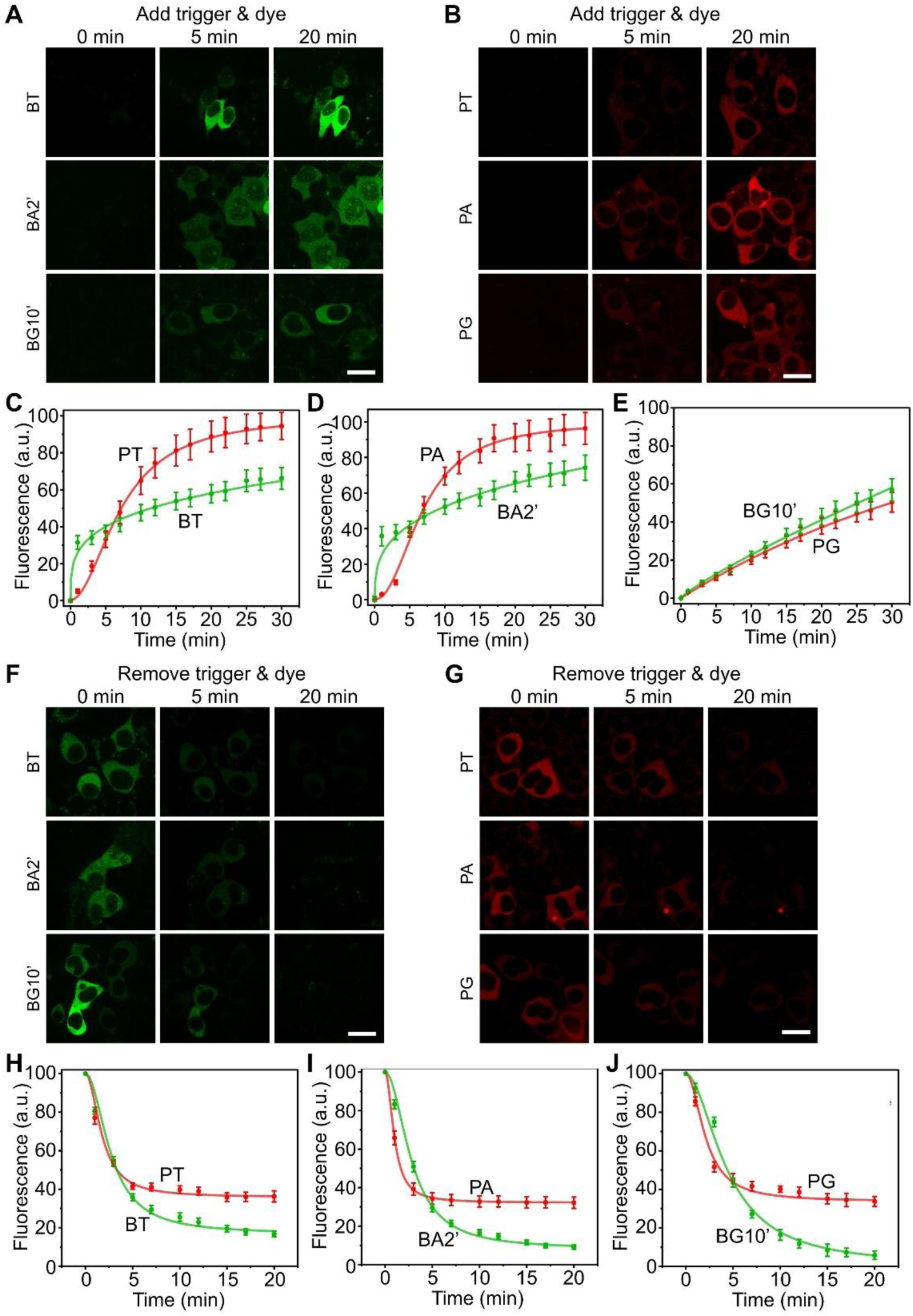
Characterization of fluorescence activation and deactivation kinetics of each trigger-activated FR reporter in living HEK293T cells. (**A**) HEK293T cells expressing trigger-activated Broccoli-based reporters were incubated with 40 μM DFHBI-1T and cognate triggers (5 μM ASP2905, 250 μM guanine, or 30 μM tetracycline), and fluorescence intensities were monitored over 30 min. Scale bar, 20 μm. (**B**) HEK293T cells expressing trigger-activated Pepper-based reporters were incubated with 1 μM HBC620 and cognate triggers (5 μM ASP2905, 250 μM guanine, or 30 μM tetracycline), and fluorescence intensities were monitored over 30 min. Scale bar, 20 μm. (**C–E**) Quantification of cellular fluorescence activation kinetics as performed by monitoring ∼20 individual HEK293T cells per condition. Data are presented as mean ± SD from images acquired in at least three independent experiments. (**F**, **G**) Deactivation kinetics were evaluated after incubating HEK293T cells expressing trigger-activated FR reporters with the corresponding dye and trigger combinations for ∼45 min to achieve maximal fluorescence activation. Cells were then imaged during signal removal following removal of both dye and trigger, followed by three sequential DPBS washes (1 min each) to initiate fluorescence decay. Scale bar, 20 μm. (**H–J**) Quantification of cellular fluorescence deactivation kinetics measured from ∼20 individual HEK293T cells per condition. Data are presented as mean ± SD from images acquired in at least three independent experiments.

Pepper-based reporters showed a similar trend (Figure 3B). Upon co-incubation with HBC620 and tetracycline, ASP2905, or guanine, ∼50% of the maximal fluorescence signal was reached within ∼7 min, ∼6 min, and ∼30 min, for the PT, PA, and PG reporters, respectively (Figure 3C–E). Although guanine-responsive reporters again activated more slowly than those responsive to tetracycline or ASP2905, their high brightness enabled readily detectable fluorescence within ∼10 min for the Pepper channels, supporting the practical time resolution of the system.

We then measured deactivation kinetics following removal of both dye and trigger. Cells were first incubated with the cognate dye-trigger combinations for ∼45 min to achieve near-maximal activation, followed by three rapid washes (1 min each) with dye-free Dulbecco’s phosphate-buffered saline (DPBS) to initiate signal decay. For Broccoli-based reporters, cellular fluorescence decreased rapidly, dropping to below 20% of the initial signal within ∼10 min (Figure 3F,H–J). In contrast, Pepper-based reporters exhibited slower deactivation. Under the same wash conditions, approximately 39%, 32%, and 40% of the initial fluorescence remained at 10 min for tetracycline-, ASP2905-, and guanine-responsive Pepper constructs, respectively, with residual signals detectable for up to ∼20 min (Figure 3G–J). These results indicate that the Pepper-based constructs require additional wash steps with longer incubation times to achieve more complete signal removal. Accordingly, in the subsequent sequential imaging experiments, five consecutive 2-min washes were implemented. Overall, these kinetic measurements indicate that the six trigger-activated reporters support rapid activation and practical signal removal, consistent with the requirements for sequential multiplexed imaging in living cells.

### Multi-cassette vector design and eight-channel live-cell sequential imaging

To enable concurrent expression of six trigger-activated fluorogenic RNAs together with two additional orthogonal FR reporters (Corn and DNB), we constructed AIO-Puro-based multi-cassette plasmids [16]. Two vectors were designed to distribute reporter modules across separate RNA scaffolds and thereby reduce undesired RNA–RNA interactions. In the first construct (**BAPG-BTPA**), circular BA2’ and PG were positioned on two arms of a three-way junction within an F30 scaffold, while circular BT and PA were incorporated into a second F30 scaffold within the same vector (Figure S4A). The F30 scaffold is known to promote proper folding of RNA aptamers while reducing unwanted interactions between neighboring RNA elements, thereby improving overall reporter performance. In the second construct (**BGC-PTD**), circular BG10’ and Corn, together with circular PT and DNB, were assembled in an analogous architecture within a separate expression cassette (Figure S4B). Different FRs and trigger combinations were intentionally distributed across distinct F30 scaffolds to minimize potential cross-reactivity and structural interference.

We next evaluated sequential imaging of trigger-activated reporters together with Corn/DFHO and DNB/TMR-DN in HEK293T cells. We implemented a dual-channel, four-round imaging workflow. In Round 1, Corn/DFHO and DNB/TMR-DN were imaged simultaneously using their well-separated excitation/emission profiles. Following the seqFRIES protocol, dyes were incubated for ∼5 min and fluorescence was subsequently removed through five consecutive 2-min DPBS washes (Figure 4A). In Rounds 2–4, trigger-activated imaging was performed sequentially: tetracycline (5-min incubation), ASP2905 (5-min incubation), and guanine (10-min incubation to ensure bright signals), each followed by five 2-min DPBS washes for signal removal. Robust fluorescence switching was observed throughout the sequential imaging and washing cycles.

**Figure 4.**
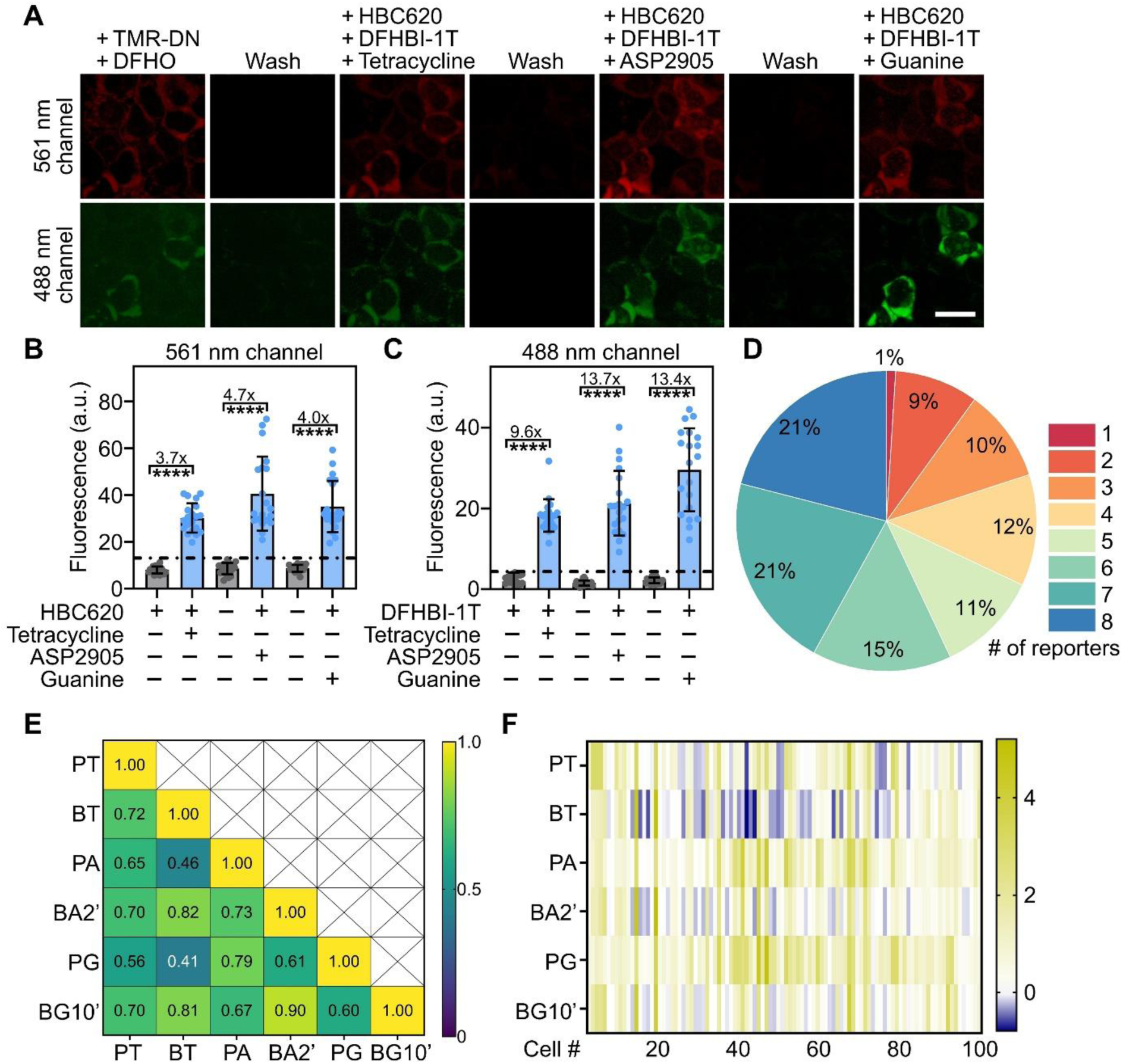
Eight-plex sequential imaging of logicFRIES in living HEK293T cells. (**A**) Sequential imaging was performed using a four-round workflow. In the first round, 10 μM DFHO and 0.5 μM TMR-DN were added for 5 min before imaging, followed by five consecutive DPBS washes (2 min each) to remove residual fluorescence signals. In the subsequent rounds, trigger-dependent imaging was performed sequentially using 30 μM tetracycline (5 min incubation), 5 μM ASP2905 (5 min incubation), and 250 μM guanine (10 min incubation), together with 40 μM DFHBI-1T and 1 μM HBC620, followed by identical washing steps as in the first round. Scale bar, 20 μm. (**B**, **C**) Quantification of signal changes during sequential imaging and washing cycles. Significant fluorescence activation and removal between imaging rounds were confirmed by one-way ANOVA analysis (****p < 0.0001) across ∼20 individual cells. (**D**) Percentage of HEK293T cells exhibiting 1–8 activated fluorescence channels following four-round sequential imaging. Data were obtained from 100 cells across at least three independent experiments. (**E**) Pearson correlation heatmap showing fluorescence correlation among the six trigger-activated FR channels measured from 100 individual HEK293T cells. Pseudocolor values represent Pearson’s correlation coefficients (*r*) between channel pairs. (**F**) Fluorescence intensity heatmap of the six trigger-activated FR channels across 100 individual HEK293T cells. Pseudocolor values represent normalized fluorescence intensities calculated by subtracting the cellular fluorescence threshold and dividing by the standard deviation of cellular fluorescence. Positive values indicate activated (“ON”) fluorescence states.

Significant fluorescence changes between imaging rounds were confirmed by fold enhancement and one-way ANOVA analysis (Figure 4B,C). For the Pepper-based reporters, fluorescence increased by ∼3.7-, 4.7-, and 4.0-fold following the addition of tetracycline, ASP2905, and guanine, respectively, relative to the preceding wash step. Similarly, the Broccoli-based reporters exhibited ∼9.6-, 13.7-, and 13.4-fold fluorescence enhancements upon the addition of tetracycline, ASP2905, and guanine, respectively. To distinguish positive fluorescence activation from background, threshold values were defined as the mean background fluorescence (μ) plus three standard deviations (σ) (i.e., μ + 3σ) using the wash step with the highest fluorescence intensity (often the final wash before ligand addition). The resulting thresholds were ∼4.3 for the Pepper reporters and ∼13.1 for the Broccoli reporters, as indicated by the dashed lines in Figure 4B,C.

Across the population, ∼57% of transfected HEK293T cells exhibited bright fluorescence across 6–8 reporter channels (Figure 4D). Trigger-activated Broccoli and Pepper signals showed strong cell-to-cell correlation, with most Pearson’s correlation coefficients (*r*) ranging from ∼0.6 to ∼0.9 (Figure 4E). PA and PG reporters were detected in nearly 100% of transfected cells (Figure 4F), whereas BT exhibited lower activation efficiency (∼60% of cells). These data together support effective activation and sequential switching of multiple logic reporters, while also revealing construct-dependent variability that likely reflects differences in expression-dependent sensitivity, RNA folding efficiency, and/or intracellular accessibility, highlighting opportunities for further optimization.

### Further validation of logicFRIES in living cells

To confirm trigger specificity and rule out cross-activation during sequential imaging, we performed control experiments in which HEK293T cells expressing either BAPG-BTPA or BGC-PTD plasmid alone were subjected to the complete imaging and stripping workflow. As expected, addition of a given trigger/dye pair in the absence of its corresponding logic reporter produced minimal fluorescence. Specifically, in BAPG-BTPA-expressing cells, co-incubation with DFHO and TMR-DN yielded negligible signal (Figure 5A), consistent with the absence of the corresponding Corn and DNB reporters in this construct (Figure 5B). Upon addition of ASP2905, both the Broccoli and Pepper channels were activated, whereas addition of tetracycline or guanine selectively activated only the relevant Broccoli or Pepper channel expected to be present in BAPG-BTPA (Figure 5C,D). Accordingly, the fluorescence signals associated with the absent reporters remained below the threshold line, confirming the lack of activation for the corresponding reporter systems.

**Figure 5.**
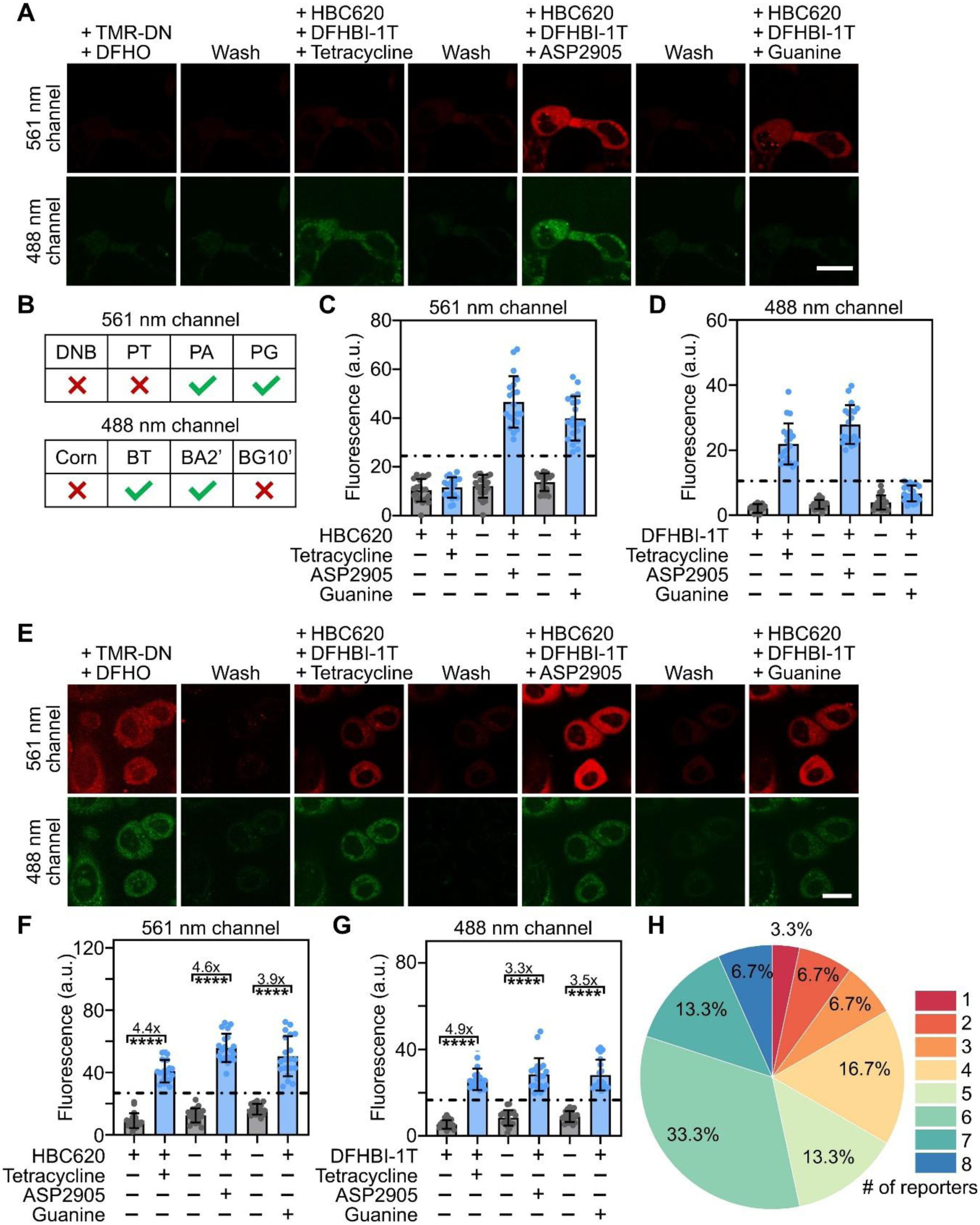
Further validation of the logicFRIES platform inside cells. (**A**) Sequential imaging was performed in HEK293T cells expressing only the BAPG-BTPA plasmid using a four-round workflow. In the first round, 20 μM DFHO and 0.5 μM TMR-DN were added for 5 min before imaging, followed by five consecutive DPBS washes (2 min each) to remove residual fluorescence signals. In the subsequent rounds, trigger-dependent imaging was performed sequentially using 30 μM tetracycline (5 min incubation), 5 μM ASP2905 (5 min incubation), and 250 μM guanine (10 min incubation), together with 40 μM DFHBI-1T and 1 μM HBC620, followed by identical washing steps as in the first round. Scale bar, 20 μm. (**B**) Presence and absence of the corresponding logicFRIES reporter within this platform. (**C**, **D**) Quantification of signal changes during sequential imaging and washing cycles across ∼20 individual cells. Data are presented as mean ± SD from at least three independent experiments. (**E**) Sequential imaging was performed in SKBR3 cells expressing both BAPG-BTPA and BGC-PTD plasmids using a four-round workflow. In the first round, 10 μM DFHO and 0.5 μM TMR-DN were added for 5 min before imaging, followed by five consecutive DPBS washes (2 min each) to remove residual fluorescence signals. In the subsequent rounds, trigger-dependent imaging was performed sequentially using 30 μM tetracycline (5 min incubation), 5 μM ASP2905 (5 min incubation), and 250 μM guanine (10 min incubation), together with 40 μM DFHBI-1T and 1 μM HBC620, followed by identical washing steps as in the first round. Scale bar, 20 μm. (**F**, **G**) Quantification of signal changes during sequential imaging and washing cycles. Significant fluorescence activation and removal between imaging rounds were confirmed by one-way ANOVA analysis (****p < 0.0001) across ∼20 individual cells. (**H**) Percentage of SKBR3 cells exhibiting 1–8 activated fluorescence channels following four-round sequential imaging. Data were obtained from 30 cells across at least three independent experiments.

Conversely, in BGC-PTD-expressing cells, minimal activation was observed following ASP2905 treatment in both Broccoli and Pepper channels, and trigger addition in the tetracycline and guanine rounds produced signals only in the channels corresponding to reporters encoded by this construct (Figure S5A,B). Quantitative analysis further confirmed that fluorescence levels were markedly reduced when the corresponding logic reporter was absent, compared with cells expressing the complete reporter set (Figure S5C,D). Together, these results demonstrate specific trigger-mediated activation and minimal cross-activation among orthogonal FR/trigger modules during sequential imaging.

We next evaluated the performance of the logicFRIES system in SKBR3 cells. As shown in Figure S6A, we first characterized each logic reporter individually by imaging cells in separate wells in the presence of its cognate trigger and dye. All FR/trigger/dye combinations produced robust fluorescence signals, confirming effective activation. Quantitative analysis further revealed significant differences between dye-only and trigger-activated conditions for both Pepper- and Broccoli-based reporters (Figure S6B,C). Specifically, the Pepper and Broccoli reporters exhibited ∼2.9–5.0-fold and ∼4.2–5.8-fold trigger-induced fluorescence activations, respectively.

We then performed eight-channel sequential imaging in SKBR3 cells using the same imaging and stripping workflow established in HEK293T cells (Figure 5E). Fluorescence intensities changed significantly after each imaging and washing step, indicating efficient signal activation and removal (Figure 5F,G). Pepper-based reporters displayed fluorescence increases of ∼4.4-fold for PT, ∼4.6-fold for PA, and ∼3.9-fold for PG compared to the previous washing step. Broccoli-based reporters showed fluorescence enhancements of ∼4.9-fold, ∼3.3-fold, and ∼3.5-fold following the addition of tetracycline, ASP2905, and guanine, respectively. Similar to HEK293T cells, a cutoff value of μ + 3σ was calculated based on the background fluorescence during the wash steps for both the Pepper and Broccoli reporters, as represented by the dashed lines in Figure 5F,G. Consistent with the HEK293T results, ∼53% of transfected SKBR3 cells exhibited detectable fluorescence from 6–8 logic reporter channels (Figure 5H). PG and PA signals were detected in nearly all transfected cells, whereas BG10’ exhibited the lowest activation efficiency, with detectable signals in ∼46% of SKBR3 cells (Figure S7A).

Correlation analysis across trigger-activated FR channels yielded moderate Pearson’s correlation coefficients, mostly ranging from ∼0.3 to ∼0.8 (Figure S7B). The observed variations in correlation values and heatmap patterns likely reflect differences in sampling depth and cell-to-cell heterogeneity in reporter expression and/or activation efficiency. Taken together, these results establish a significantly enhanced multiplexing platform capable of visualizing at least eight intracellular targets in living mammalian cells within ∼1 hour. This workflow supports rapid, sequential, and specific live-cell imaging, with strong potential for dynamic and high-content cellular analysis.

## Conclusion

In this study, to the best of our knowledge, we demonstrate the highest-order multiplexed live-cell RNA imaging reported to date. Eight fluorogenic RNA-based reporters are integrated, and an optimized sequential imaging and stripping workflow is established. By leveraging the reversible interaction between fluorogenic RNAs and their cognate dyes and triggers, this approach mitigates spectral constraints inherent to conventional fluorescence imaging and enables repeated visualization of multiple molecular signals within the same living cells. This logicFRIES system exhibited robust trigger-dependent fluorescence activation with minimal cross-reactivity among orthogonal triggers. Sequential imaging in HEK293T and SKBR3 cells confirmed that multiple reporters can be visualized within individual cells, with the majority of transfected cells displaying signals across multiple channels. Furthermore, specificity controls demonstrated that fluorescence activation depended strongly on the presence of both the appropriate reporter and its cognate trigger, confirming the orthogonality and reliability of the expanded reporter system.

Beyond fluorescence tagging, the modular and programmable nature of fluorogenic RNA reporters creates a clear path toward multiplexed molecular sensing. As a future direction, logic AND-gated FR sensors could be engineered to detect endogenous RNAs, analogous to prior seqFRIES designs [16]. In this architecture, a blocker domain hybridizes to the FR in the absence of the target RNA, preventing dye-pocket formation and minimizing background (Figure S8). Only concurrent target RNA and trigger binding induces the active FR conformation and restores fluorescence, adding specificity for selective imaging of endogenous RNAs in heterogeneous cell populations.

Similarly, the programmability of FR reporters could be extended to proteins, metabolites, and signaling molecules by incorporating target-specific aptamer sequences into other critical stems of the reporter architecture. Simultaneous binding of the target analyte and trigger molecule would drive conformational rearrangement, restore the dye-binding pocket, and generate fluorescence. Such modular strategies could transform logicFRIES into a broadly applicable platform for multiplexed detection of various classes of biomolecule targets.

Overall, this work establishes the feasibility of an expanded sequential imaging platform for multiplexed reporting in living systems. Continued development of orthogonal trigger sets and FR/dye pairs, together with advances in automated imaging and fluidic control, should further enable logicFRIES for more comprehensive characterization of complex cellular states. These tools can be potentially applied to resolve relationships among multiple biomarkers and to quantify cell-to-cell heterogeneity, enabling advanced live-cell molecular profiling.

## Acknowledgements

The authors gratefully acknowledge the support from NSF 2435059, Chan Zuckerberg Initiative Dynamic Imaging program 2023-321170, and Camille Dreyfus Teacher-Scholar Award to M.Y. The authors also thank other members of the You Lab for useful discussion and valuable comments.

## Data Availability Statement

Experimental data that supports the findings of this study is available in the Supporting Information.

---

**Scheme 1.**
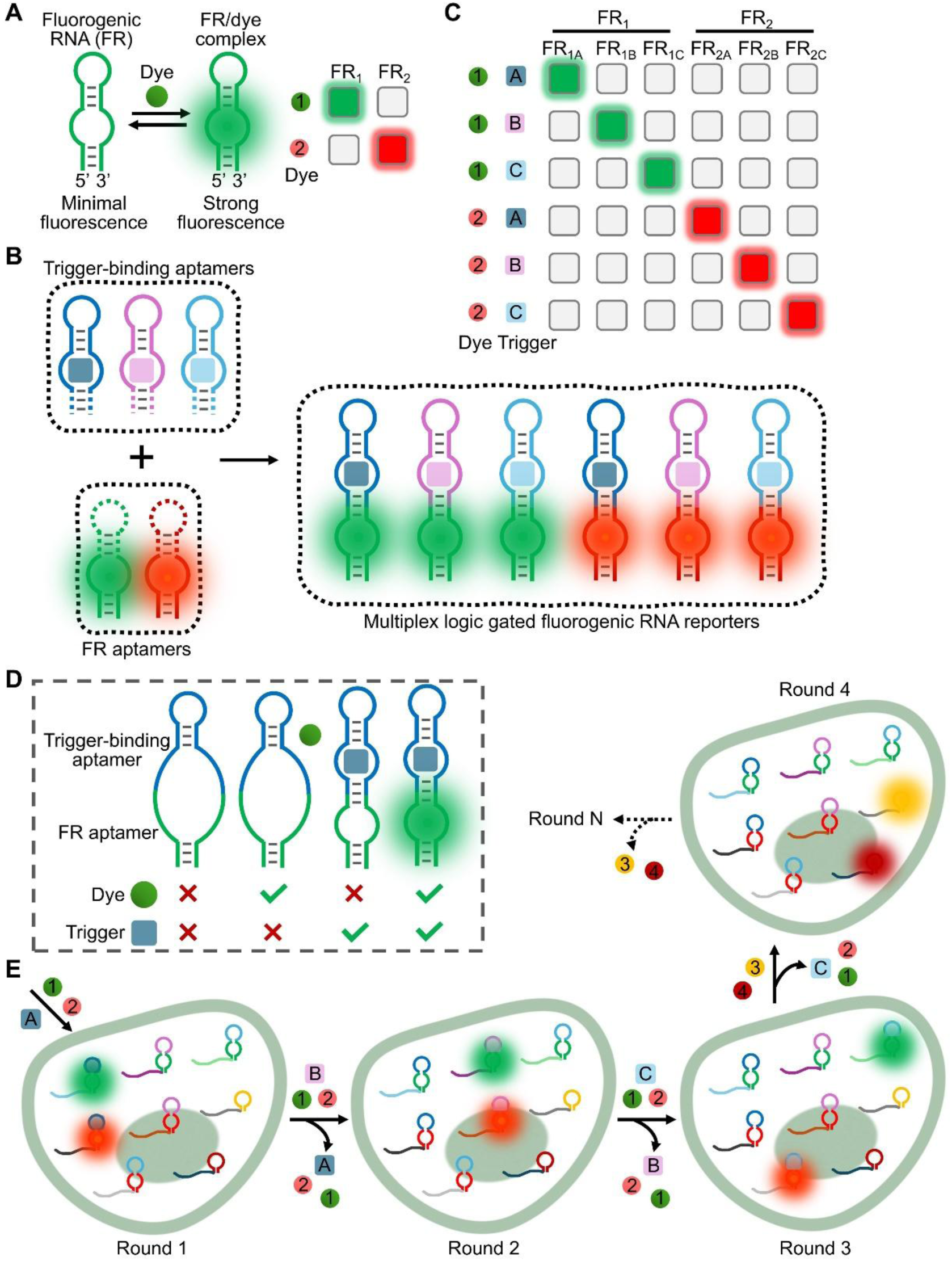
Working principle of fluorogenic RNA (FR) aptamers and logicFRIES. (**A**) FR aptamers selectively bind otherwise nonfluorescent small-molecule dyes and activate fluorescence; orthogonal FR/dye pairs enable multiplexed imaging. (**B**, **C**) LogicFRIES expands multiplexing by fusing FR reporters with trigger-binding RNA aptamers to generate trigger-responsive, AND-gated reporters in which fluorescence requires two independent inputs, i.e., trigger binding and dye binding, yielding three trigger-defined activation states per FR and six distinguishable signals from two FRs. (**D**) Schematic of the four conformational states of an AND-gated reporter in the presence or absence of the cognate trigger and dye. (**E**) Overall logicFRIES workflow: six logic-gated reporters are combined with two additional orthogonal FR/dye pairs. Sequential addition of trigger and dye pairs, imaging, and wash-based signal stripping enables eight-plex live-cell imaging in four imaging rounds.

