## Supplementary Information for "Logic-Gated Fluorogenic RNA Reporters for Multiplexed Live-Cell Imaging"

† These authors contribute equally.

* Corresponding author

### Supplementary Methods

**Reagents.** All chemicals were obtained from MilliporeSigma or Fisher Scientific unless otherwise specified and were used without further purification. The compound 4-(2-hydroxyethyl-methylamino)-benzylidene-cyanophenylacetonitrile analog (HBC620) and ASP2905 were purchased from GlpBio (cat. GC60186 & GC61960). 3,5-Difluoro-4-hydroxybenzylidene imidazolinone-2-oxime (DFHO) was obtained from Bio-Techne (cat. 6434), and tetramethylrhodamine-2,4-dinitroaniline (TMR-DN) was purchased from Lumiprobe (cat. 2641).

**Synthesis of oligonucleotides and general methods.** Single-stranded DNA oligonucleotides were purchased from Integrated DNA Technologies (Coralville, IA) or the W. M. Keck Oligonucleotide Synthesis Facility (Yale University School of Medicine) and were cartridge-purified. These oligonucleotides were dissolved in syringe-filtered TE buffer (10 mM Tris-HCl, 0.1 mM EDTA, pH 7.5) to a final concentration of 100 μM and stored at −20 °C. Double-stranded DNA fragments (gBlocks) were obtained from Integrated DNA Technologies and reconstituted in TE buffer according to the manufacturer's instructions. DNA templates and inserts were generated by PCR amplification using an Eppendorf Mastercycler and Q5 High-Fidelity 2× Master Mix (New England Biolabs, M0492). PCR products were purified using the Monarch PCR & DNA Cleanup Kit (NEB, T1030). DNA concentrations were determined using a NanoDrop One UV-Vis spectrophotometer. PCR products were further analyzed by 3% (w/v) TBE agarose gel electrophoresis for approximately 30 min. RNA was synthesized via *in vitro* transcription using the HiScribe™ T7 High Yield RNA Synthesis Kit (NEB, E2040S). Following DNase I (RNase-free) treatment to remove DNA templates, RNA transcripts were purified using column-based methods and verified by 10% denaturing polyacrylamide gel electrophoresis. Purified RNA samples were aliquoted and stored at −20 °C for short-term use or at −80 °C for long-term storage. RNA secondary structures were designed and analyzed using the NUPACK and UNAFold online platforms.

***In vitro* fluorescence assay.** Fluorescence measurements were carried out at room temperature (23°C) using a PTI fluorimeter (Horiba, NJ). Samples were prepared in a buffer containing 40 mM Tris, 5 mM $MgCl_2$, and 100 mM KCl (pH 7.6). Broccoli and Pepper fluorescence spectra were recorded with excitation wavelengths at 472 nm and 577 nm, respectively. All fluorescence data were processed and plotted using the GraphPad Prism software.

**Vector construction for mammalian cell imaging.** Plasmids encoding circular RNA scaffolds were constructed using the pAV-U6+27-Tornado vector (Addgene, 124360), which includes a U6 promoter and self-cleaving ribozymes that enable autocatalytic RNA circularization. The vector backbone was prepared by double digestion with NotI-HF (NEB, R3189) and SacII (NEB, R0157). DNA inserts were generated either by PCR amplification or obtained as gBlocks from Integrated DNA Technologies, each designed with 25-bp overlaps for Gibson assembly. Assembly reactions were performed using Gibson Assembly Master Mix (NEB, E2611) according to the manufacturer's instructions. For constructs containing multiple RNA scaffolds, BAPG-BTPA and BGC-PTD were cloned into the AIO-Puro vector (Addgene, 74630). Inserts were digested with BsaI-HFv2 (NEB, R3733S) or BbsI-HF (NEB, R3539S) and ligated into the corresponding restriction sites using T4 DNA ligase (NEB, B0202S). All ligation products were transformed into NEB 5α chemically competent *E. coli* (NEB, C2987) and selected based on ampicillin resistance. Plasmids were extracted using the GeneJET Plasmid Miniprep Kit (Thermo Scientific, FERK0503). Sequence verification was performed by Sanger sequencing (Eurofins Genomics) or whole-plasmid sequencing (Azenta Life Sciences).

**Mammalian cell culture and transfection.** HEK293T/17 (CRL-11268) and SKBR3 cell lines were obtained from ATCC (American Type Culture Collection). Cells were cultured in Dulbecco's Modified

Eagle Medium (Thermo Scientific, 11995-065) and McCoy's 5A Medium (Thermo Scientific,16-600-108) supplemented with 10% fetal bovine serum (Thermo Scientific, A5670801) and 1% penicillin-streptomycin at 37 °C in a 5% $CO_2$ atmosphere. All cell lines were confirmed to be free of mycoplasma contamination. Cells were passaged at approximately 80% confluence using TrypLE Express (Thermo Scientific, 12604013). For HEK293T transfection, FuGENE HD (Promega, E2312) was used following the manufacturer's protocol. Briefly, 2.5 µg total plasmid DNA (1:1 ratio for co-transfections) was mixed with 100 µL Opti-MEM (Thermo Scientific, 31985062) and 7 µL FuGENE HD, incubated for 15 min at room temperature, and 12 µL of the mixture was added to each well of an 8-well chamber slide (Cellvis, C8-1.5P). Untransfected cells were used as negative controls. For SKBR3 cells, transfection was performed using Lipofectamine™ 3000 (Invitrogen). Two mixtures were prepared: Tube 1 containing 25 µL Opti-MEM and 0.5 µL Lipofectamine 3000, and Tube 2 containing 25 µL Opti-MEM, 500 ng DNA, and 1 µL P3000 reagent. After combining and incubating for 10–15 min, 25 µL of the complex was added per well. Cells were imaged 48 hours post-transfection.

**Mammalian cell imaging and data analysis.** Live-cell imaging was performed using a Yokogawa spinning disk confocal system mounted on a Nikon Eclipse Ti inverted microscope and controlled by NIS-Elements AR software. Images were acquired using a 40× oil immersion objective. Broccoli/DFHBI-1T and Corn/DFHO were excited at 488 nm, with emission collected between 500–550 nm. Pepper/HBC620 and DNB/TMR-DN were excited at 561 nm, with emission collected between 575–625 nm. Cytosolic regions were manually selected for quantitative analysis using ImageJ. Data processing and curve fitting were performed using Origin and GraphPad Prism. Background fluorescence was determined using untransfected HEK293T and SKBR3 cells incubated for 30 min with triggers (30 µM tetracycline, 10 µM ASP2905, or 250 µM guanine) and dyes at the following concentrations: 40 µM DFHBI-1T, 1 µM HBC620, 0.5 µM TMR-DN, or 40 µM DFHO. Threshold values were calculated as the mean background fluorescence ($\mu$) plus three standard deviations ($\sigma$), defined as $\mu + 3\sigma$. Cells exhibiting fluorescence intensities above this threshold were classified as "ON", while those below were considered "OFF".

### Supplementary Figures

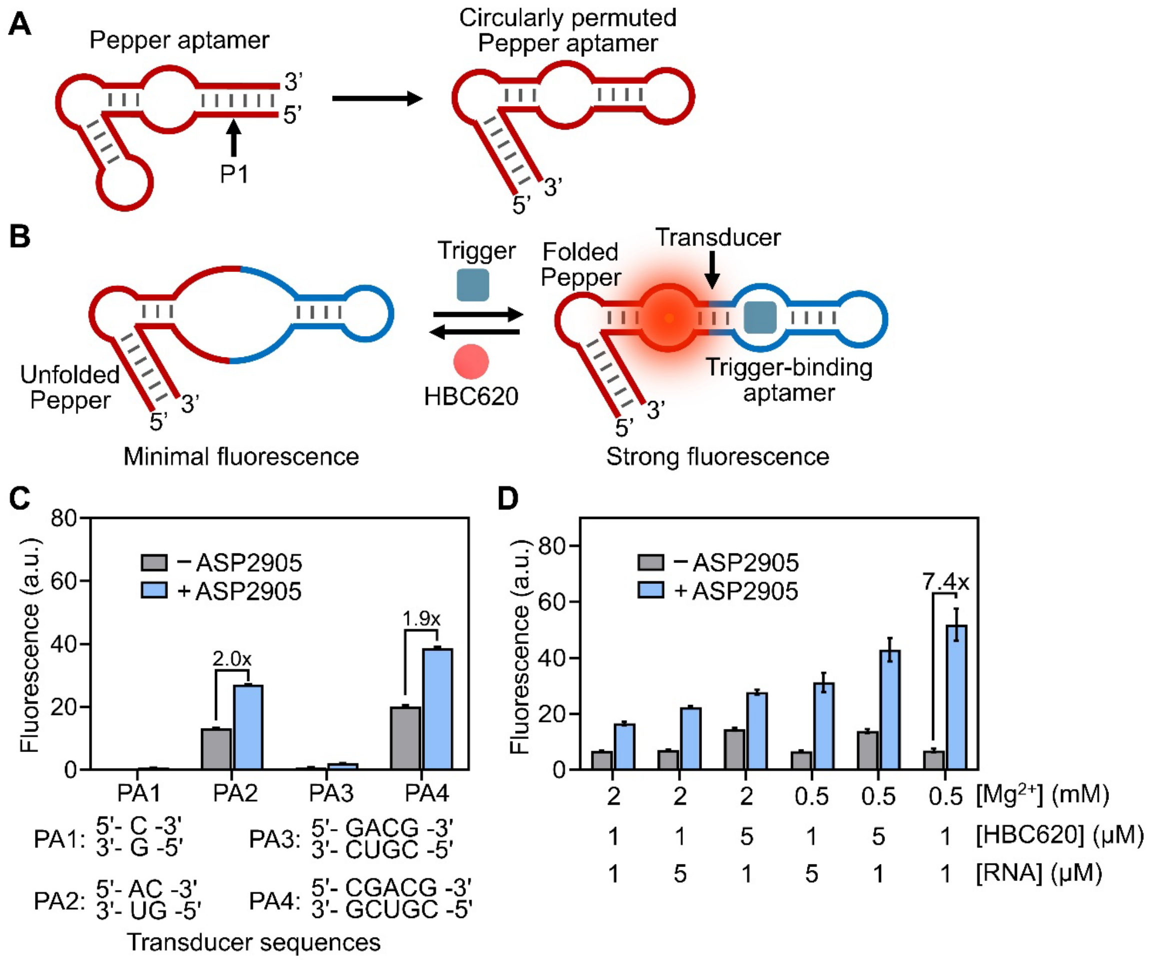


**Figure S1.** Design and *in vitro* optimization of trigger-activated Pepper reporters. (**A**) Schematic of the Pepper aptamer and the circularly permuted Pepper construct. The highlighted P1 stem is a critical structural element required for proper folding and fluorescence. (**B**) Design of the trigger-activated circularly permuted Pepper reporter, in which a trigger-binding aptamer and transducer domain are integrated into the Pepper P1 stem. In the absence of trigger, the transducer disfavors formation of the active Pepper fold. Trigger binding stabilizes the aptamer, promotes transducer strand hybridization, and restores the native Pepper conformation, thereby activating fluorescence. (**C**) Trigger-dependent fluorescence activation of ASP2905-responsive Pepper reporter variants (PA1–PA4) and their corresponding transducer sequences. Fluorescence was measured after incubating 1 μM RNA and 5 μM HBC620 dye in the presence or absence of 5 μM ASP2905. (**D**) *In vitro* optimization of the ASP2905-responsive Pepper reporter PA2. Fluorescence was measured as a function of $Mg^{2+}$, HBC620 dye, and RNA concentrations. All data are reported as mean ± standard deviation (SD) from at least three independent experiments and are normalized to the fluorescence of the Pepper/HBC620 positive control measured under matched conditions.

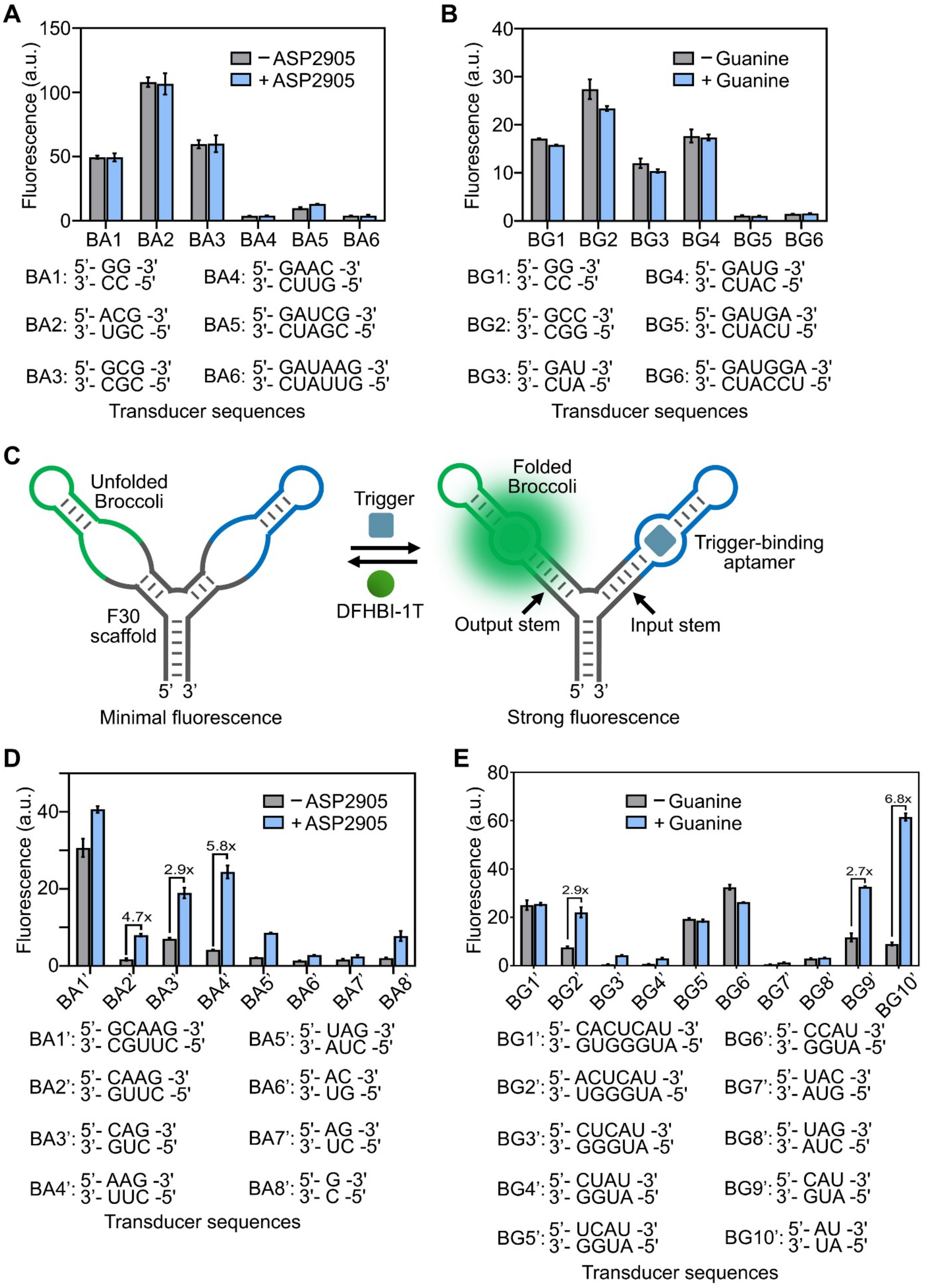


**Figure S2.** Design and *in vitro* characterization of trigger-activated Broccoli reporters. (**A**) Trigger-dependent fluorescence activation of initial allosteric ASP2905-responsive Broccoli reporter variants

(BA1–BA6) and their corresponding transducer sequences. Fluorescence was measured after incubating 1 μM RNA and 20 μM DFHBI-1T dye in the presence or absence of 5 μM ASP2905. (**B**) Trigger-dependent fluorescence activation of initial allosteric guanine-responsive Broccoli reporter variants (BG1–BG6) and their corresponding transducer sequences. Fluorescence was measured after incubating 1 μM RNA and 20 μM DFHBI-1T dye in the presence or absence of 100 μM guanine. (**C**) Schematic of an alternative RNA nanodevice framework using an F30 RNA scaffold to replace the conventional duplex transducer, providing a three-way junction architecture that supports ligand-dependent control of Broccoli folding. Trigger binding induces a conformational rearrangement of the trigger aptamer that stabilizes the input stem, which subsequently promotes folding of the output stem. This structural transition enables Broccoli to bind DFHBI-1T and generate a fluorescence signal. (**D**) Trigger-dependent fluorescence activation of alternative three-way junction ASP2905-responsive Broccoli reporter variants (BA1'–BA8') and their corresponding transducer sequences. Fluorescence was measured after incubating 1 μM RNA and 20 μM DFHBI-1T dye in the presence or absence of 5 μM ASP2905. (**B**) Trigger-dependent fluorescence activation of alternative three-way junction guanine-responsive Broccoli reporter variants (BG1'–BG10') and their corresponding transducer sequences. Fluorescence was measured after incubating 1 μM RNA and 20 μM DFHBI-1T dye in the presence or absence of 100 μM guanine. All data are reported as mean ± SD from at least three independent experiments and are normalized to the fluorescence of the Broccoli/DFHBI-1T positive control measured under matched conditions.

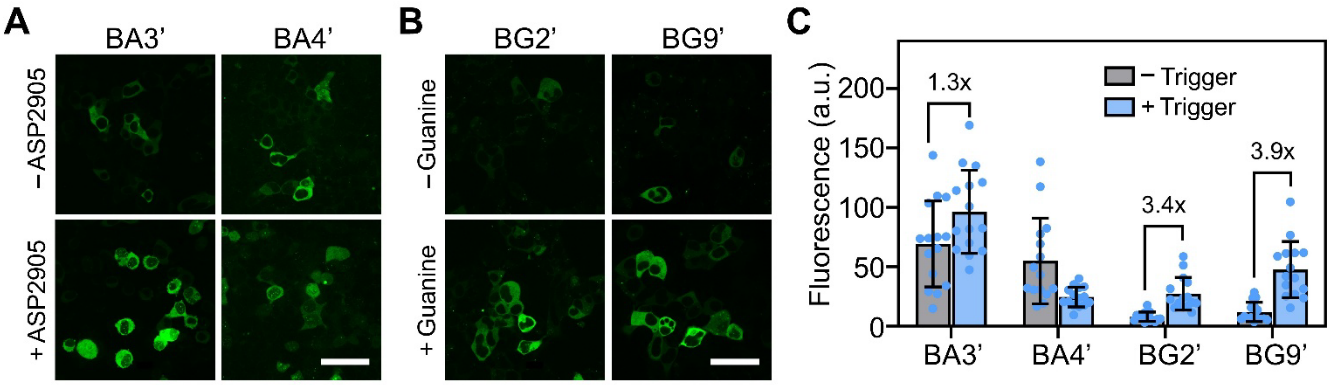


**Figure S3.** Optimization and evaluation of trigger-activated Broccoli reporters in living cells. (**A**) Fluorescence imaging of HEK293T cells expressing circular ASP2905-responsive Broccoli reporter variants from the pAV-U6+27 vector after incubation with 40 μM DFHBI-1T dye in the presence or absence of 5 μM ASP2905. Scale bar, 50 μm. (**B**) Fluorescence imaging of HEK293T cells expressing circular guanine-responsive Broccoli reporter variants from the pAV-U6+27 vector after incubation with 40 μM DFHBI-1T dye in the presence or absence of 250 μM guanine. Scale bar, 50 μm. (**C**) Intracellular trigger-activated fluorescence intensities were quantified from ~20 individual cells per condition. Data are presented as mean ± SD from images acquired in at least three independent experiments and normalized to Broccoli positive controls after incubation with 40 μM DFHBI-1T.

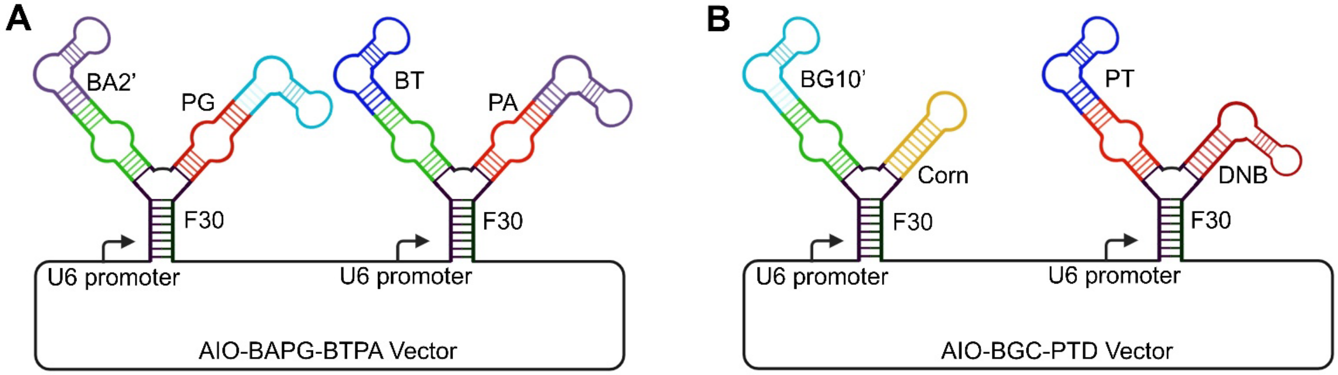


**Figure S4.** Design of multi-cassette AIO-BAPG-BTPA and AIO-BGC-PTD plasmids. (**A**) Schematic illustration of the AIO-BAPG-BTPA construct, in which circular BA2' and PG were positioned on two arms of a three-way junction within an F30 scaffold, while circular BT and PA were incorporated into a second F30 scaffold within the same vector. (**B**) Schematic illustration of the AIO-BGC-PTD construct containing circular BG10' and Corn, together with circular PT and DNB, arranged on separate F30 scaffolds.

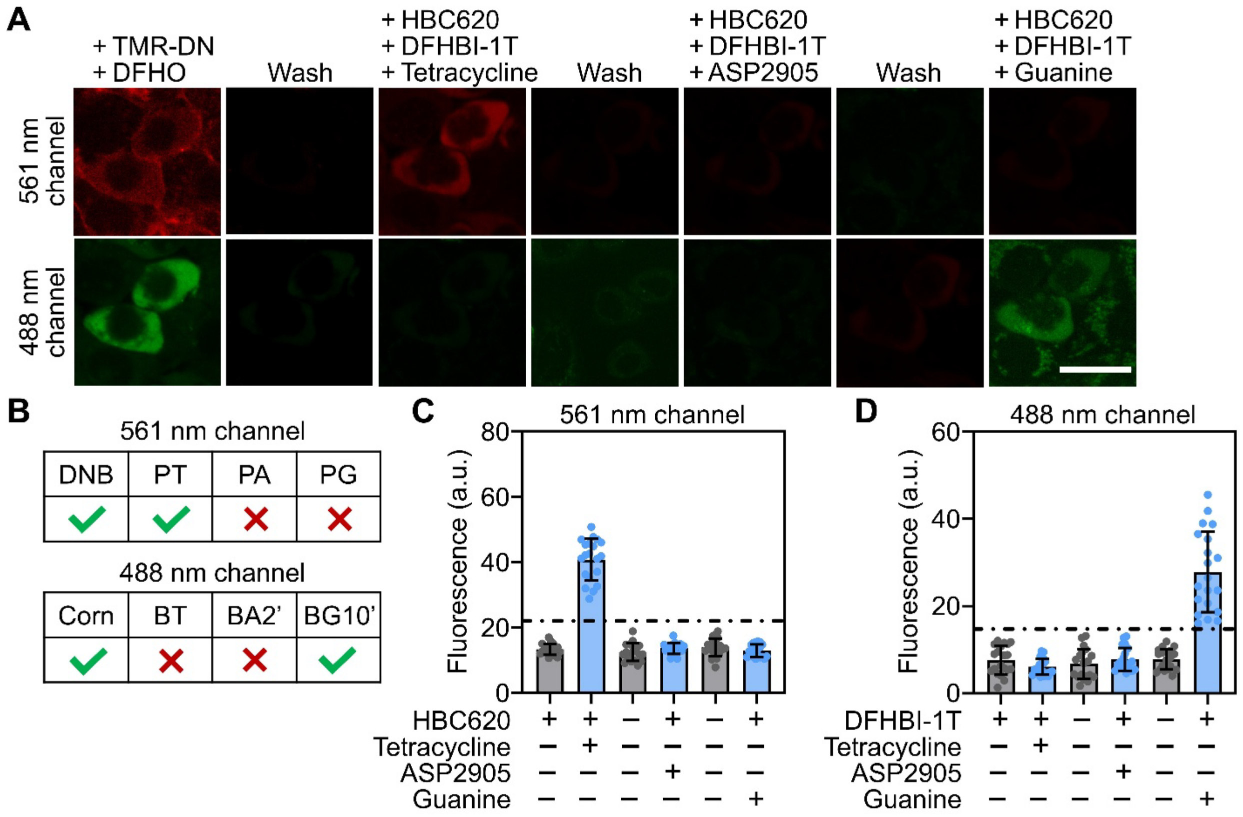


**Figure S5.** Further validation of the logicFRIES platform inside cells. (**A**) Sequential imaging was performed in HEK293T cells expressing only the BGC-PTD plasmid using a four-round workflow. In the first round, 20 μM DFHO and 0.5 μM TMR-DN were added for 5 min before imaging, followed by five consecutive DPBS washes (2 min each) to remove residual fluorescence signals. In the subsequent rounds, trigger-dependent imaging was performed sequentially using 30 μM tetracycline (5 min incubation), 5 μM ASP2905 (5 min incubation), and 250 μM guanine (10 min incubation), together with 40 μM DFHBI-1T and 1 μM HBC620, followed by identical washing steps as in the first round. Scale bar, 20 μm. (**B**) Presence and absence of the corresponding logicFRIES reporter within this platform. (**C**, **D**) Quantification of signal changes during sequential imaging and washing cycles across ~20 individual cells. Data are presented as mean ± SD from at least three independent experiments.

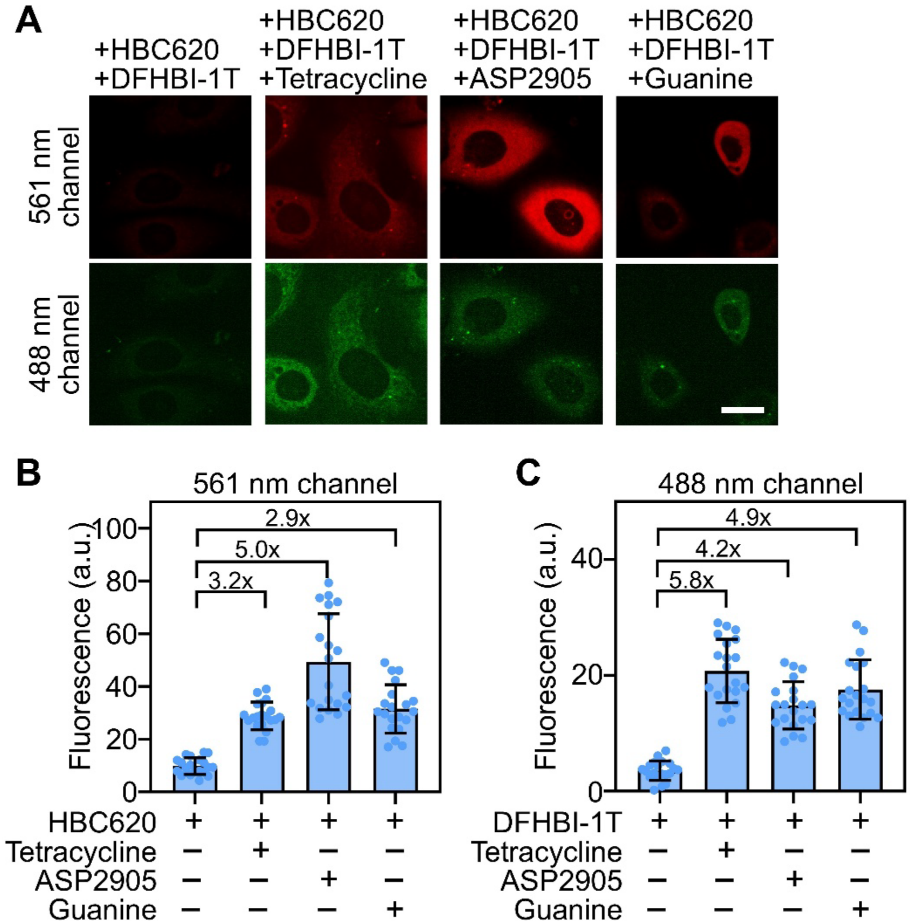


**Figure S6.** Characterization of logicFRIES reporters in SKBR3 cells. (**A**) Fluorescence imaging of SKBR3 cells expressing both AIO-BAPG-BTPA and AIO-BGC-PTD plasmids after incubation with 40 μM DFHBI-1T dye and 1 μM HBC620 in the presence or absence of individual triggers, including 30 μM tetracycline, 5 μM ASP2905, and 250 μM guanine. Scale bar, 20 μm. (**B**) Quantification of trigger-activated FR reporters in response to cognate and non-cognate triggers, measured from ~20 individual cells per condition. Data are presented as mean ± SD from images acquired in at least three independent experiments and normalized to the corresponding Pepper/HBC620 positive controls. (**C**) Quantification of trigger-activated FR reporters in response to cognate and non-cognate triggers, measured from ~20 individual cells per condition. Data are presented as mean ± SD from images acquired in at least three independent experiments and normalized to the corresponding Broccoli/DFHBI-1T positive controls.

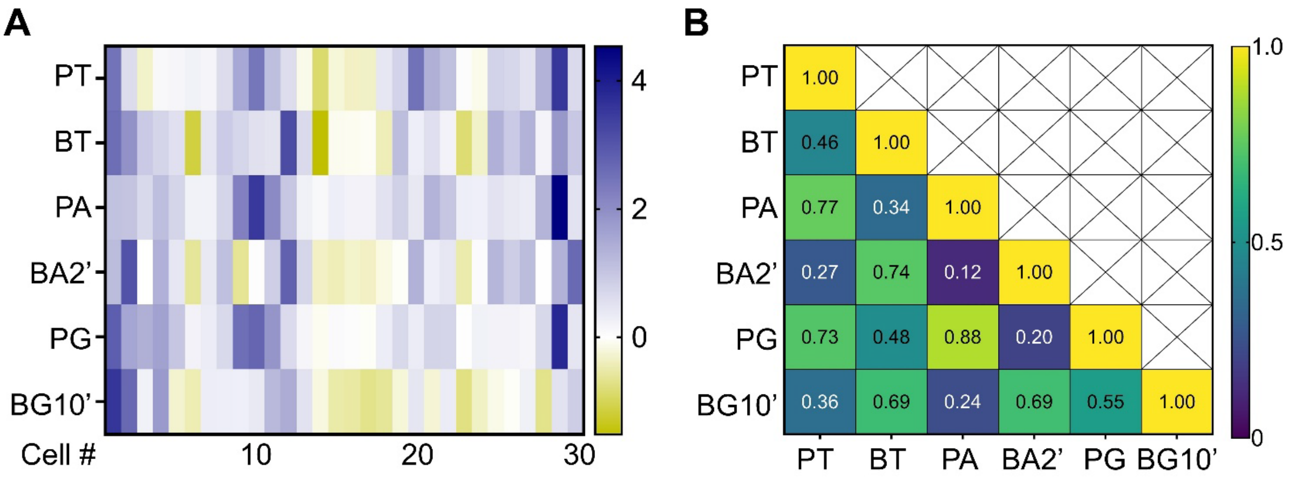


**Figure S7.** Sequential multiplexed analysis with logicFRIES in SKBR3 cells. (**A**) Fluorescence intensity heatmap of the six trigger-activated FR channels across 30 individual SKBR3 cells. Pseudocolor values represent normalized fluorescence intensities calculated by subtracting the cellular fluorescence threshold and dividing by the standard deviation of the cellular fluorescence signal. Positive values indicate activated ("ON") fluorescence states. (**B**) Pearson correlation heatmap showing fluorescence correlations among the six trigger-activated FR channels across 30 individual SKBR3 cells. Pseudocolor values represent Pearson's correlation coefficients ($r$) calculated for each pair of fluorescence channels.

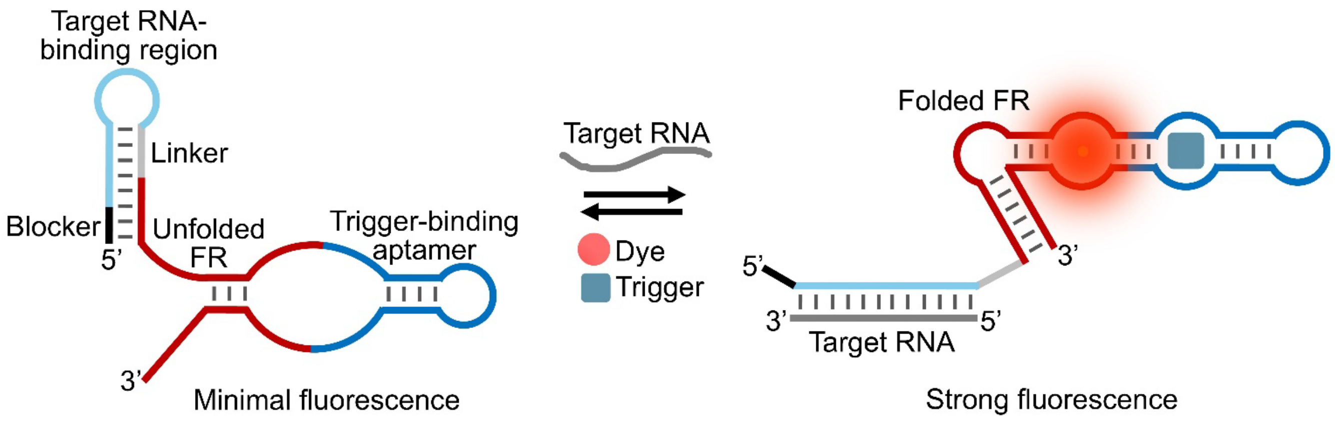


**Figure S8.** Schematic illustration of converting a logicFRIES reporter into a sensor for detecting endogenous RNA targets. The sensor is composed of a blocker, target-binding region, linker, and trigger-activated FR. In the absence of either the target RNA or the cognate trigger, the FR remains misfolded and nonfluorescent. Simultaneous binding of the target RNA and trigger induces a structural rearrangement of the sensor, restoring the active FR conformation and enabling fluorescence, thereby providing highly specific detection of endogenous RNA targets with minimal background signal.

### Supplementary Table

**Table S1.** List of RNA sequences used in this work. **Bolded** nucleotides are sequences of fluorogenic RNAs (Broccoli, Pepper, Corn, or DNB). Blue nucleotides indicate sequences for the F30 scaffold (**Bolded blue** is the F30 scaffold in the AIO-Puro plasmid). Red nucleotides illustrate trigger-specific aptamers. Underlined nucleotides represent the transducer. Purple nucleotides are ribozyme cutting sites for further RNA cyclization. Green and orange nucleotides show input and output stems, respectively.

| Name | RNA sequence (5' to 3') |
|---|---|
| PA1 | **GGGUCCAAUCGUGGCGUGUCG**CUGAGAGAGACGGAUUCCGUCCGCGAAUUCACGCUGG**ACUGGCGCCGACCC** |
| PA2 | **GGGUCCAAUCGUGGCGUGUCG**ACUGAGAGAGACGGAUUCCGUCCGCGAAUUCACGCUGGGU**ACUGGCGCCGACCC** |
| PA3 | **GGGUCCAAUCGUGGCGUGUCG**GACGUGAGAGAGACGGAUUCCGUCCGCGAAUUCACGCUGCGUC**ACUGGCGCCGACCC** |
| PA4 | **GGGUCCAAUCGUGGCGUGUCG**CGACGUGAGAGAGACGGAUUCCGUCCGCGAAUUCACGCUGCGUCG**ACUGGCGCCGACCC** |
| PT | **GGCCACCCAAUCGUGGCGUGUCG**AUAAAACAUACCAGAUUUCGAUCUGGAGAGGUGAAGAAUACGACCACCUAU**ACUGGCGCCGGUGGCC** |
| PG | **GGCCACCCAAUCGUGGCGUGUCG**CAUAUAAUCGCGUGGAUAUGGCACGCAAGUUUCUACCGGGCACCGUAAAUGUCCGACUAUG**ACUGGCGCCGGUGGCC** |
| BA1 | **GCGGAGACGGUCGGGUCCA**GGUGAGAGAGACGGAUUCCGUCCGCGAAUUCACGCUGCC**UGUCGAGUAGAGUGUGGGCUCCGC** |
| BA2 | **GCGGAGACGGUCGGGUCCA**ACGUGAGAGAGACGGAUUCCGUCCGCGAAUUCACGCUGCGU**UGUCGAGUAGAGUGUGGGCUCCGC** |
| BA3 | **GCGGAGACGGUCGGGUCCA**GCGUGAGAGAGACGGAUUCCGUCCGCGAAUUCACGCUGCGC**UGUCGAGUAGAGUGUGGGCUCCGC** |
| BA4 | **GCGGAGACGGUCGGGUCCA**GAACUGAGAGAGACGGAUUCCGUCCGCGAAUUCACGCUGGUUC**UGUCGAGUAGAGUGUGGGCUCCGC** |
| BA5 | **GCGGAGACGGUCGGGUCCA**GAUCGUGAGAGAGACGGAUUCCGUCCGCGAAUUCACGCUGCGAUC**UGUCGAGUAGAGUGUGGGCUCCGC** |
| BA6 | **GCGGAGACGGUCGGGUCCA**GAUAAGUGAGAGAGACGGAUUCCGUCCGCGAAUUCACGCUGGUUAUC**UGUCGAGUAGAGUGUGGGCUCCGC** |
| BG1 | **GCGGAGACGGUCGGGUCCA**GGAUAAUCGCGUGGAUAUGGCACGCAAGUUUCUACCGGGCACCGUAAAUGUCCGACUCC**UGUCGAGUAGAGUGUGGGCUCCGC** |
| BG2 | **GCGGAGACGGUCGGGUCCA**GCCAUAAUCGCGUGGAUAUGGCACGCAAGUUUCUACCGGGCACCGUAAAUGUCCGACUGGC**UGUCGAGUAGAGUGUGGGCUCCGC** |
| BG3 | **GCGGAGACGGUCGGGUCCA**GAUAUAAUCGCGUGGAUAUGGCACGCAAGUUUCUACCGGGCACCGUAAAUGUCCGACUAUC**UGUCGAGUAGAGUGUGGGCUCCGC** |
| BG4 | **GCGGAGACGGUCGGGUCCA**GAUGAUAAUCGCGUGGAUAUGGCACGCAAGUUUCUACCGGGCACCGUAAAUGUCCGACUCAUC**UGUCGAGUAGAGUGUGGGCUCCGC** |

| | |
|---|---|
| BG5 | **GCGGAGACGGUCGGGUCCA**GAUGAAUAAUCGCGUGGAUAUGGCACGCAAGUUUCUACCGGGCACCGUAAAUGUCCGACUUCAUC**UGUCGAGUAGAGUGUGGGCUCCGC** |
| BG6 | **GCGGAGACGGUCGGGUCCA**GAUGGAAUAAUCGCGUGGAUAUGGCACGCAAGUUUCUACCGGGCACCGUAAAUGUCCGACUUCCAUC**UGUCGAGUAGAGUGUGGGCUCCGC** |
| BA1’ | GGUUGCCAUGUGUUCU**GUCGAGUAGAGUGUGGGCUCUUCGGAGACGGUCGGGUCC**AGAACUCUGGCAAGUGAGAGAGACGGAUUCCGUCCGCGAAUUCACGCUGCUUGCUCAUGGCAACC |
| BA2’ | GGUUGCCAUGUGUUCU**GUCGAGUAGAGUGUGGGCUCUUCGGAGACGGUCGGGUCC**AGAACUCUGCAAGUGAGAGAGACGGAUUCCGUCCGCGAAUUCACGCUGCUUGUCAUGGCAACC |
| BA3’ | GGUUGCCAUGUGUUCU**GUCGAGUAGAGUGUGGGCUCUUCGGAGACGGUCGGGUCC**AGAACUCUGCAGUGAGAGAGACGGAUUCCGUCCGCGAAUUCACGCUGCUGUCAUGGCAACC |
| BA4’ | GGUUGCCAUGUGUUCU**GUCGAGUAGAGUGUGGGCUCUUCGGAGACGGUCGGGUCC**AGAACUCUGAAGUGAGAGAGACGGAUUCCGUCCGCGAAUUCACGCUGCUUUCAUGGCAACC |
| BA5’ | GGUUGCCAUGUGUUCU**GUCGAGUAGAGUGUGGGCUCUUCGGAGACGGUCGGGUCC**AGAACUCUGUAGUGAGAGAGACGGAUUCCGUCCGCGAAUUCACGCUGCUAUCAUGGCAACC |
| BA6’ | GGUUGCCAUGUGUUCU**GUCGAGUAGAGUGUGGGCUCUUCGGAGACGGUCGGGUCC**AGAACUCUGACUGAGAGAGACGGAUUCCGUCCGCGAAUUCACGCUGGUUCAUGGCAACC |
| BA7’ | GGUUGCCAUGUGUUCU**GUCGAGUAGAGUGUGGGCUCUUCGGAGACGGUCGGGUCC**AGAACUCUGAGUGAGAGAGACGGAUUCCGUCCGCGAAUUCACGCUGCUUCAUGGCAACC |
| BA8’ | GGUUGCCAUGUGUUCU**GUCGAGUAGAGUGUGGGCUCUUCGGAGACGGUCGGGUCC**AGAACUCUGGUGAGAGAGACGGAUUCCGUCCGCGAAUUCACGCUGCUCAUGGCAACC |
| BG1’ | GGUUGCCAUGUGUUCU**GUCGAGUAGAGUGUGGGCUCUUCGGAGACGGUCGGGUCC**AGAACUCUGCACUCAUAUAAUCGCGUGGAUAUGGCACGCAAGUUUCUACCGGGCACCGUAAAUGUCCGACUAUGGGUGUCAUGGCAACC |
| BG2’ | GGUUGCCAUGUGUUCU**GUCGAGUAGAGUGUGGGCUCUUCGGAGACGGUCGGGUCC**AGAACUCUGACUCAUAUAAUCGCGUGGAUAUGGCACGCAAGUUUCUACCGGGCACCGUAAAUGUCCGACUAUGGGUUCAUGGCAACC |
| BG3’ | GGUUGCCAUGUGUUCU**GUCGAGUAGAGUGUGGGCUCUUCGGAGACGGUCGGGUCC**AGAACUCUGCUCAUAUAAUCGCGUGGAUAUGGCACGCAAGUUUCUACCGGGCACCGUAAAUGUCCGACUAUGGGUCAUGGCAACC |
| BG4’ | GGUUGCCAUGUGUUCU**GUCGAGUAGAGUGUGGGCUCUUCGGAGACGGUCGGGUCC**AGAACUCUGCUAUAUAAUCGCGUGGAUAUGGCACGCAAGUUUCUACCGGGCACCGUAAAUGUCCGACUAUGGUCAUGGCAACC |

| | |
|---|---|
| BG5' | GGUUGCCAUGUGUUCU**GUCGAGUAGAGUGUGGGCUCUUCGGAGACGGUCGGGUC** **C**AGAACUCUGUCAUAUAAUCGCGUGGAUAUGGCACGCAAGUUUCUACCGGGCACCG UAAAUGUCCGACUAUGGUCAUGGCAACC |
| BG6' | GGUUGCCAUGUGUUCU**GUCGAGUAGAGUGUGGGCUCUUCGGAGACGGUCGGGUC** **C**AGAACUCUGCCAUAUAAUCGCGUGGAUAUGGCACGCAAGUUUCUACCGGGCACCG UAAAUGUCCGACUAUGGUCAUGGCAACC |
| BG7' | GGUUGCCAUGUGUUCU**GUCGAGUAGAGUGUGGGCUCUUCGGAGACGGUCGGGUC** **C**AGAACUCUGUACAUAAUCGCGUGGAUAUGGCACGCAAGUUUCUACCGGGCACCGU AAAUGUCCGACUGUAUCAUGGCAACC |
| BG8' | GGUUGCCAUGUGUUCU**GUCGAGUAGAGUGUGGGCUCUUCGGAGACGGUCGGGUC** **C**AGAACUCUGUAGAUAAUCGCGUGGAUAUGGCACGCAAGUUUCUACCGGGCACCGU AAAUGUCCGACUCUAUCAUGGCAACC |
| BG9' | GGUUGCCAUGUGUUCU**GUCGAGUAGAGUGUGGGCUCUUCGGAGACGGUCGGGUC** **C**AGAACUCUGCAUAUAAUCGCGUGGAUAUGGCACGCAAGUUUCUACCGGGCACCGU AAAUGUCCGACUAUGUCAUGGCAACC |
| BG10' | GGUUGCCAUGUGUUCU**GUCGAGUAGAGUGUGGGCUCUUCGGAGACGGUCGGGUC** **C**AGAACUCUGAUAUAAUCGCGUGGAUAUGGCACGCAAGUUUCUACCGGGCACCGUA AAUGUCCGACUAUUCAUGGCAACC |
| Cyclic PT | GGCCGCACUCGCCGGUCCCAAGCCCGGAUAAAAUGGGAGGGGGCGGGAAACCGCC UAACCAUGCCGAGUGCGGCCGC**GGCCACCCAAUCGUGGCGUGUCG**<u>AU</u>AAAACAUAC CAGAUUUCGAUCUGGAGAGGUGAAGAAUACGACCACCU<u>AU</u>**ACUGGCGCCGGUGGCC** GUGGCCGCGGUCGGCGUGGACUGUAGAACACUGCCAAUGCCGGUCCCAAGCCCGG AUAAAA GUGGAGGGUACAGUCCACGC |
| Cyclic PA | GGCCGCACUCGCCGGUCCCAAGCCCGGAUAAAAUGGGAGGGGGCGGGAAACCGCC UAACCAUGCCGAGUGCGGCCGC**GGGUCCAAUCGUGGCGUGUCG**<u>AC</u>UGAGAGAGAC GGAUUCCGUCCGCGAAUUCACGCUG<u>GU</u>**ACUGGCGCCGACCC**GUGGCCGCGGUCGG CGUGGACUGUAGAACACUGCCAAUGCCGGUCCCAAGCCCGGAUAAAA GUGGAGGGUACAGUCCACGC |
| Cyclic PG | GGCCGCACUCGCCGGUCCCAAGCCCGGAUAAAAUGGGAGGGGGCGGGAAACCGCC UAACCAUGCCGAGUGCGGCCGC**GGCCACCCAAUCGUGGCGUGUCG**<u>CAU</u>AUAAUCG CGUGGAUAUGGCACGCAAGUUUCUACCGGGCACCGUAAAUGUCCGACU<u>AUG</u>**ACUGG** **CGCCGGUGGCC**GUGGCCGCGGUCGGCGUGGACUGUAGAACACUGCCAAUGCCGGU CCCAAGCCCGGAUAAAA GUGGAGGGUACAGUCCACGC |
| Cyclic BT | GGCCGCACUCGCCGGUCCCAAGCCCGGAUAAAAUGGGAGGGGGCGGGAAACCGCC UAACCAUGCCGAGUGCGGCCGC**GCGGAGACGGUCGGGUCCAG**<u>AUGGA</u>AAAACAUA CCAGAUUUCGAUCUGGAGAGGUGAAGAAUACGACCACCU<u>UCCCA</u>**CUGUCGAGUAGA** **GUGUGGGCUCCGC**GUGGCCGCGGUCGGCGUGGACUGUAGAACACUGCCAAUGCCG GUCCCAAGCCCGGAUAAAA GUGGAGGGUACAGUCCACGC |
| Cyclic BA2' | GGCCGCACUCGCCGGUCCCAAGCCCGGAUAAAAUGGGAGGGGGCGGGAAACCGCC UAACCAUGCCGAGUGCGGCCGCGGUUGCCAUGUGUUCU**GUCGAGUAGAGUGUGGG** **CUCUUCGGAGACGGUCGGGUCC**AGAACUCUGCAAGUGAGAGAGACGGAUUCCGUC CGCGAAUUCACGCUGCUUGUCAUGGCAACCGUGGCCGCGGUCGGCGUGGACUGUA GAACACUGCCAAUGCCGGUCCCAAGCCCGGAUAAAA GUGGAGGGUACAGUCCACGC |

| | |
|---|---|
| Cyclic BA3’ | GGCCGCACUCGCCGGUCCCAAGCCCGGAUAAAAUGGAGGGGGCGGGAAACCGCCUAACCAUGCCGAGUGCGGCCGCGGUUGCCAUGUGUUCU**GUCGAGUAGAGUGUGGGCUCUUCGGAGACGGUCGGGUCC**AGAACUCUGCAGUGAGAGAGACGGAUUCCGUCCGCGAAUUCACGCUGCUGUCAUGGCAACCGUGGCCGCGGUCGGCGUGGACUGUAGAACACUGCCAAUGCCGGUCCCAAGCCCGGAUAAAA GUGGAGGGUACAGUCCACGC |
| Cyclic BA4’ | GGCCGCACUCGCCGGUCCCAAGCCCGGAUAAAAUGGAGGGGGCGGGAAACCGCCUAACCAUGCCGAGUGCGGCCGCGGUUGCCAUGUGUUCU**GUCGAGUAGAGUGUGGGCUCUUCGGAGACGGUCGGGUCC**AGAACUCUGAAGUGAGAGAGACGGAUUCCGUCCGCGAAUUCACGCUGCUUUCAUGGCAACCGUGGCCGCGGUCGGCGUGGACUGUAGAACACUGCCAAUGCCGGUCCCAAGCCCGGAUAAAA GUGGAGGGUACAGUCCACGC |
| Cyclic BG2’ | GGCCGCACUCGCCGGUCCCAAGCCCGGAUAAAAUGGAGGGGGCGGGAAACCGCCUAACCAUGCCGAGUGCGGCCGCGGUUGCCAUGUGUUCU**GUCGAGUAGAGUGUGGGCUCUUCGGAGACGGUCGGGUCC**AGAACUCUGACUCAUAUAAUCGCGUGGAUAUGGCACGCAAGUUUCUACCGGGCACCGUAAAUGUCCGACUAUGGGUCAUGGCAACCGUGGCCGCGGUCGGCGUGGACUGUAGAACACUGCCAAUGCCGGUCCCAAGCCCGGAUAAAA GUGGAGGGUACAGUCCACGC |
| Cyclic BG9’ | GGCCGCACUCGCCGGUCCCAAGCCCGGAUAAAAUGGAGGGGGCGGGAAACCGCCUAACCAUGCCGAGUGCGGCCGCGGUUGCCAUGUGUUCU**GUCGAGUAGAGUGUGGGCUCUUCGGAGACGGUCGGGUCC**AGAACUCUGCAUAUAAUCGCGUGGAUAUGGCACGCAAGUUUCUACCGGGCACCGUAAAUGUCCGACUAUGUCAUGGCAACCGUGGCCGCGGUCGGCGUGGACUGUAGAACACUGCCAAUGCCGGUCCCAAGCCCGGAUAAAAGUGGAGGGUACAGUCCACGC |
| Cyclic BG10’ | GGCCGCACUCGCCGGUCCCAAGCCCGGAUAAAAUGGAGGGGGCGGGAAACCGCCUAACCAUGCCGAGUGCGGCCGCGGUUGCCAUGUGUUCU**GUCGAGUAGAGUGUGGGCUCUUCGGAGACGGUCGGGUCC**AGAACUCUGAUAUAAUCGCGUGGAUAUGGCACGCAAGUUUCUACCGGGCACCGUAAAUGUCCGACUAUUCAUGGCAACCGUGGCCGCGGUCGGCGUGGACUGUAGAACACUGCCAAUGCCGGUCCCAAGCCCGGAUAAAAGUGGAGGGUACAGUCCACGC |
| BAPG | GCCGC**UUGCCAUGUGUAUCG**GGUUGCCAUGUGUUCU**GUCGAGUAGAGUGUGGGCUCUUCGGAGACGGUCGGGUCC**AGAACUCUGCAAGUGAGAGAGACGGAUUCCGUCCGCGAAUUCACGCUGCUUGUCAUGGCAACC**CGAUACUCUGAUGAUCCGGCCACCCAAUCGUGGCGUGUCG**CAUAUAAUCGCGUGGAUAUGGCACGCAAGUUUCUACCGGGCACCGUAAAUGUCCGACUAUG**ACUGGCGCCGGUGGCCGGAUCAUUCAUGGCAA** |
| BTPA | GCCGC**UUGCCAUGUGUAUCGGCGGAGACGGUCGGGUCCAG**AUGGAAAAACAUACCAGAUUUCGAUCUGGAGAGGUGAAGAAUACGACCACCUUCCCA**CUGUCGAGUAGAGUGUGGGCUCCGCCGAUACUCUGAUGAUCCGGGUCCAAUCGUGGCGUGUCG**ACUGAGAGAGACGGAUUCCGUCCGCGAAUUCACGCUGGU**ACUGGCGCCGACCCGGAUCAUUCAUGGCAA** |
| BGC | GCCGC**UUGCCAUGUGUAUCG**GGUUGCCAUGUGUUCU**GUCGAGUAGAGUGUGGGCUCUUCGGAGACGGUCGGGUCC**AGAACUCUGAUAUAAUCGCGUGGAUAUGGCACGCAAGUUUCUACCGGGCACCGUAAAUGUCCGACUAUUCAUGGCAACC**CGAUACUCUGAUGAUCCGGCGCGAGGAAGGAGGUCUGAGGAGGUCACUGCGCCGGAUCAUUCAUGGCAA** |

| | |
|---|---|
| PTD | GCCGC**UUGCCAUGUGUAUCGGGCCACCCAAUCGUGGCGUGUCG**AUAAAACAUACC<br>AGAUUUCGAUCUGGAGAGGUGAAGAAUACGACCACCUAU**ACUGGCGCCGGUGGCCC**<br>**GAUACUCUGAUGAUCCGGUGCCUUAUUCCGGACGCCGGGCCCGAAUGCUGCUACG**<br>**GCAGUCGAAGACAACAUCGCGCCCUUCGGAGGCACCGGAUCAUUCAUGGCAA** |
